# Epigenetic mechanisms to propagate histone acetylation by p300/CBP

**DOI:** 10.1101/2023.03.31.535039

**Authors:** Masaki Kikuchi, Satoshi Morita, Masatoshi Wakamori, Shin Sato, Tomomi Uchikubo-Kamo, Mikako Shirouzu, Takashi Umehara

## Abstract

Histone acetylation is important for the activation of gene transcription but little is known about its direct ‘read/write’ mechanisms. Here, we report cryo-electron microscopy structures in which a p300/CBP multidomain monomer recognizes histone H4 N-terminal tail (NT) acetylation (ac) in a nucleosome and acetylates non-H4 histone NTs within the same nucleosome. p300/CBP not only recognized H4NTac *via* the bromodomain pocket responsible for ‘reading’, but also interacted with the DNA minor grooves *via* the outside of that pocket. This directed the catalytic center of p300/CBP to one of the non-H4 histone NTs. The primary target that p300 ‘writes’ by ‘reading’ H4NTac was H2BNT, and H2BNTac promoted H2A-H2B dissociation from the nucleosome. We propose a model in which p300/CBP ‘replicates’ histone NT acetylation within the H3-H4 tetramer to inherit epigenetic storage, and ‘transcribes’ it from the H3-H4 tetramer to the H2B-H2A dimers to activate context-dependent gene transcription through local nucleosome destabilization.

---

In eukaryotes, genomic DNA is packaged within the cell nucleus by forming the nucleosome, the structural unit of chromatin^1^. The nucleosome, 11 nm in diameter and 5.5 nm high, consists of 145–147 base pairs (bp) of DNA wrapped around a histone octamer composed of one [H3-H4]_2_ tetramer and two [H2A-H2B] dimers^2^. In the nucleosome, eight NTs of four histone pairs protrude outside of DNA. On the side chains of each of these NTs, various post-translational modifications (PTMs) are chemically ‘written’, ‘read’, or ‘erased’ by diverse proteins^3, 4^ A typical PTM is acetylation of the Nε nitrogen of the lysine (K) side chain of histone NTs, which correlates with transcriptional activation of genes in eukaryotes^5^.

Representative ‘writers’ of lysine acetylation (Kac) are E1A-binding protein p300 (EP300; KAT3B)^6^ and its homolog, CREB-binding protein (CBP; KAT3A)^7^, which can transfer an acetyl group onto the NT lysine residue(s) for all four histones^8, 9, 10^. A typical ‘reader’ of Kac is the bromodomain, a four-helix bundle that forms a pocket preferring Kac^11^. p300 and CBP are unique in that they also contain bromodomains that can ‘read’ Kac, for example, acetylated K12 of H4 (H4K12ac)^12, 13^. Indeed, p300 acetylates NT of H2A.Z, an evolutionarily conserved variant of H2A, through the bromodomain-mediated H4NTac ‘reader’ activity^14^. Interestingly, H2BNT is acetylated only by p300/CBP^15^, which is considered a genuine signature of active enhancers and their target promoters^16^.

Although H3K27ac, a residue acetylated by p300/CBP, has been used as a marker for active promoters and enhancers^17^, it is absent in many p300-enriched chromatin regions^18^ and is dispensable for enhancer activity of gene transcription in mouse embryonic stem cells^19^. In contrast, possibly through transcription-coupled histone exchange, H2BNTac better correlates with enhancer activity than any other known chromatin marks, with RNA transcription found in 79% of all H2BNTac-positive regions^16^. Importantly, the acetyltransferase activity of p300 responsible for H2BNTac^15^ is a key driver of rapid enhancer activation and is essential for promoting the recruitment of RNA polymerase II (RNAPII) at virtually all enhancers and enhancer-regulated genes^20^. However, knowledge of how p300/CBP acetylates H2BNT and possibly thus activates transcription from enhancers and their target promoters has remained elusive.

Crystal structures of the histone acetyltransferase domain (HAT)^21^ and a multidomain encompassing bromodomain, a RING finger (RING), a plant homeodomain finger (PHD), and HAT (BRPH)^13, 22^ of p300 suggest regulatory mechanisms for the catalytic reaction of p300/CBP. Recently, cryogenic electron microscopy (cryo-EM) structure analysis of a catalytic-dead p300 multidomain complexed with the unmodified nucleosome core particle

(NCP) was reported^23^. However, the molecular mechanism of how p300/CBP ‘reads/writes’ histone acetylation in the nucleosome(s) is unknown. In this study, we report cryo-EM structures revealing how the p300/CBP multidomain involved in the ‘read/write’ of histone acetylation recognizes H4NTac containing di-acetylation at K12 and K16 (H4K12ac/K16ac) and acetylates non-H4 histone NTs in the same nucleosome.

## Results

### Dependence of histone NTac on prior H4NTac

To understand how p300/CBP ‘reads/writes’ histone acetylation, we purified catalytically active p300_BRPHZT_ (residues 1048–1836) protein, containing bromodomain, RING, and PHD (BRP), the catalytically active HAT domain with the autoinhibitory loop (AIL), ZZ, and the TAZ2 domain (BRPHZT; Fig. 1a, Supplementary Fig. 1a, b). For the nucleosome, based on the fact that p300_BRP_ preferentially binds to H4NT containing H4K12ac/K16ac^13^, we reconstituted a nucleosome, hereafter referred to as H4acNuc, consisting of the histone octamer having H4K12ac/K16ac and 146-bp palindromic human α-satellite DNA with 17-bp linker DNA linked to either end (*i.e*., a 180-bp nucleosome having H4K12ac/K16ac; Supplementary Fig. 1c, d).

**Fig. 1.**
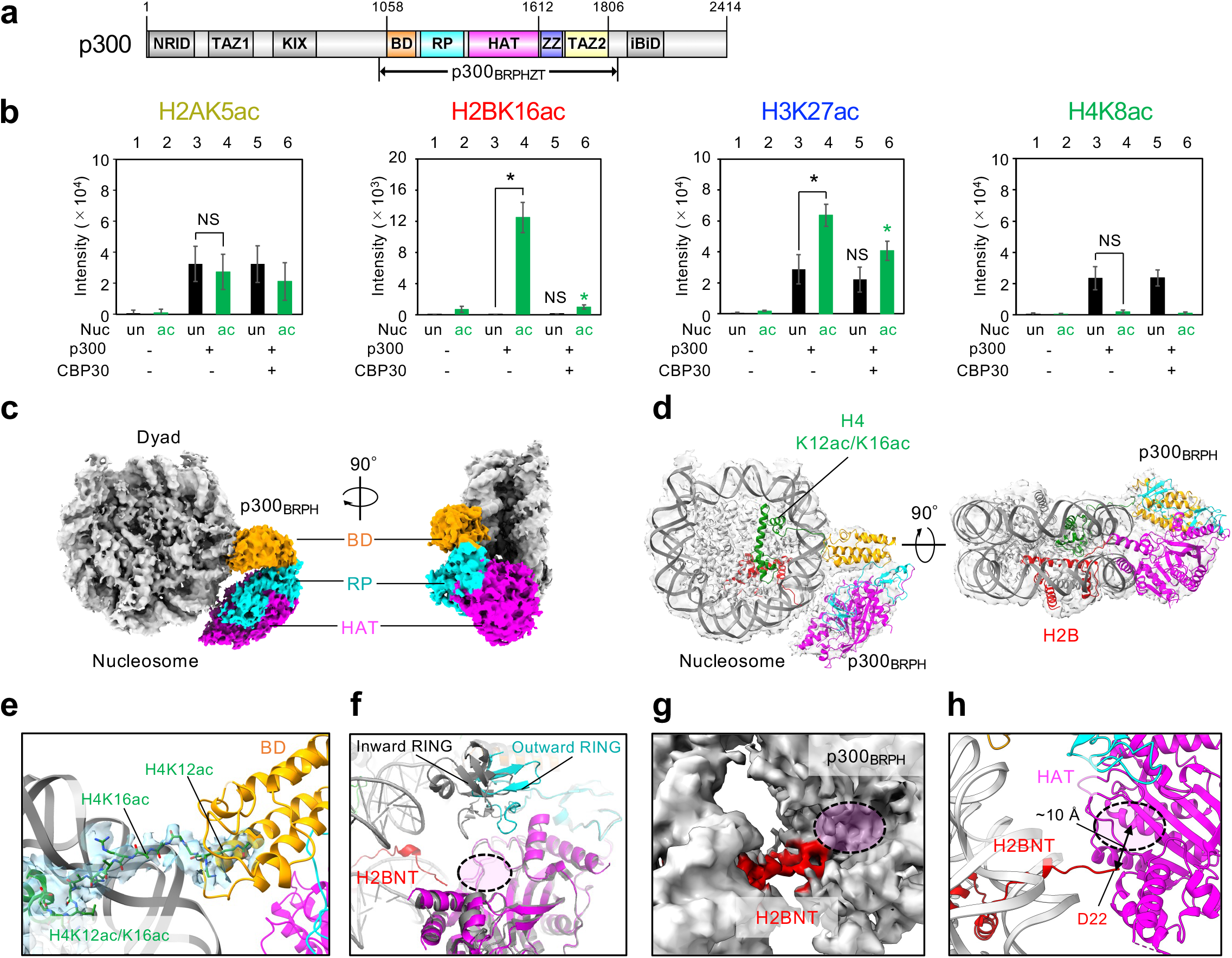
Structure of p300_BRPHZT_ bound to N-terminal tails (NT) in histone H2B and acetylated H4. **a** Schematic representation of the domain architecture of human p300. NRID, nuclear receptor interaction domain; TAZ1, transcriptional adaptor zinc-finger domain 1; KIX, kinase-inducible domain of CREB-interacting domain; BD, bromodomain; RP, the RING and PHD zinc-fingers; HAT, histone acetyltransferase domain; ZZ, ZZ-type zinc-finger; TAZ2, transcriptional adaptor zinc-finger domain 2; and IBiD, IRF3-binding domain. The positions of the N- and C-termini and the start/end residues of the major domains are shown at the top. The positions of the start/end residues of the construct used in this study (*i.e*., p300_BRPHZT_) are shown at the bottom. **b** *In vitro* acetyltransferase activity of p300_BRPHZT_ toward an H4-di-acetylated nucleosome. The histone and residue for which acetylation was detected by immunoblotting are shown above each panel. Color code: H2A, yellow; H2B, red; H3, blue; H4, green. Nucleosome (Nuc): un, unmodified; ac (green), H4K12/K16-acetylated. p300_BRPHZT_ (p300): -, none; +, 1 μM. CBP30: -, none; +, 10 μM. The y-axis indicates the immunoblotting signal intensity at 1 min after the reaction. Means ± SD (*N* = 3). Statistical significance was assessed by a two-sample one-sided Welch’s *t*-test (NS, *P* ≥ 0.05; \**P* < 0.05; \*\**P* < 0.01). The alternative hypothesis is as follows: lane 4, increase vs. lane 3; lane 5, decrease vs. lane 3; lane 6, decrease vs. lane 4. **c** Structure of p300_BRPH_ bound to H2BNT and acetylated H4NT delineated by cryo-electron microscopy (cryo-EM). Left, top view; right, side view. p300_BRPH_ (#1 in Supplementary Fig. 8) binds to H4acNuc in a Slinky-like bent conformation *via* bromodomain and HAT. **d** Overall structure of p300_H2B_ (#1) with H4-di-acetylated nucleosome in cartoon presentation. Color code: orange, p300 BD; cyan, p300 RP; magenta, p300 HAT; green, K12/K16-acetylated H4; red, H2B. **e** Close-up view of the binding mode of p300 bromodomain (BD, #1) to the H4-di-acetylated nucleosome (H4K12acK16ac). The map corresponding to H4NT is colored light blue. **f** Superposition of the cryo-EM structure of p300_BRPH_ (#4) and the crystal structure of p300_BRPH_ lacking AIL (5LKU). The magenta region circled in black is the substrate-binding site of HAT. **g** Close-up view of the cryo-EM map (#4) and the structure of H2BNT. **h** Close-up view of H2BNT (#4) shown as a cartoon representation.

First, we examined whether K12ac/K16ac in H4 facilitates p300 to acetylate the nucleosomal histone NTs and for which residues (Fig. 1b, Supplementary Fig. 2). Consistent with previous reports^8, 24^, p300_BRPHZT_ acetylated NTs of all four histones in at least one residue (*e.g*., H2AK5ac, H2BK16ac, H3K27ac, and H4K8ac), even when the nucleosome was unmodified. But overall, it acetylated NTs more rapidly in the H4acNuc than the unmodified nucleosome at the one-minute timepoint. With H4K12ac/K16ac present, p300_BRPHZT_-catalyzed acetylation increased most prominently in H2BNT, increased significantly in H3NT (except H3K18ac), and decreased at K5/K8 in H4NT (probably due to proximity caused by p300 binding to H4K12ac/K16ac). In the presence of CBP30^25^, an inhibitor that prevents the bromodomain pocket of p300 from binding to the acetylated histone NTs, the p300_BRPHZT_-catalyzed acetylation was selectively decreased at H2BK16ac and H3K27ac (Fig. 1b). These results suggest that when p300_BRPHZT_ ‘reads’ H4NTac at K12/K16, it primarily ‘writes’ Kac on H2BNT and H3NT.

### Structure of p300 bound to H4NTac and H2BNT

To elucidate the molecular mechanism by which p300/CBP ‘reads/writes’ histone acetylation of H4acNuc, we performed cryo-EM single particle structural analysis of catalytically active p300_BRPHZT_. Our preliminary experiments using a 146-bp NCP containing H4K12ac/K16ac yielded artificial structures in which p300_BRPHZT_ bound across both ends of DNA (Supplementary Fig. 3a). To avoid this unnatural binding, we used nucleosome containing linker DNA. When the 180-bp nucleosome without H4K12ac/K16ac was used, we could not determine the structure of any of the obtained classes because the density resolution corresponding to p300_BRPHZT_ was 8–12 Å (Supplementary Fig. 3b). The final three classes of p300 were each in close proximity to different histone NTs of the nucleosome and not to a specific histone NT. When the 180-bp nucleosome containing H4K12ac/K16ac (*i.e*., H4acNuc) was used, we obtained a group of structures in which p300_BRPHZT_ binds to H4acNuc in several different modes by three dimensional (3D) classification (Supplementary Fig. 4, 5). Of these complexes, the p300_BRPHZT_·H4acNuc complex, in which HAT is in close proximity to H2BNT, had the largest number of single particles and its structure could be determined at the highest resolution (p300H2BoH4acNuc; Fig. 1c, d). We obtained the 3D reconstruction density maps of p300_H2B_·H4acNuc at 3.2–4.7 Å (Supplementary Table 1). In p300_BRPHZT_, cryo-EM maps of BRPH were detected, but not of AIL, ZZ, or TAZ2. The structure-determined p300 multidomain (p300_BRPH_) bound to H4acNuc in a Slinky-like bent conformation *via* bromodomain and HAT.

The bromodomain pocket of p300_BRPH_ recognized the K12ac sidechain of H4NTac (Fig. 1e). RING of p300_BRPH_ was structured in an outward-rotated conformation^13^ (Fig. 1f), suggesting that HAT of p300_BRPH_ in p300H2B oH4acNuc is substrate-accessible. Around HAT of p300_BRPH_, the density of H2BNT toward HAT could be modeled for the C-terminal residues after D22. Density maps were also detected near the substrate-binding pocket of HAT (Fig. 1g), but the residues of H2BNT bound to HAT could not be determined. The distance between the substrate-binding pocket and D22 (~10 Å) suggests that p300_BRPH_ of this complex acetylates lysine residues from the N-terminal side of H2BNT up to K16 (Fig. 1h). Besides p300_H2B_·H4acNuc, there were several similar complexes at low resolution in which H2BNT was located near the substrate-binding pocket of HAT (Supplementary Fig. 4, 5). Thus, it is likely that HAT of p300_BRPH_ can acetylate various lysine residues around K16 of H2BNT by a similar mechanism. These results provide a structural basis for a ‘read/write’ mechanism by which p300 recognizes H4NTac and acetylates H2BNT within the same nucleosome.

### Multiple modes of binding to histone NTs

Consistent with the fact that K12ac/K16ac in H4NT facilitated the p300_BRPHZT_-catalyzed acetylation of multiple non-H4 histone NTs (Fig. 1b, Supplementary Fig. 2), we obtained several cryo-EM structure classes in which HAT of P300_BRPHZT_ bound with non-H4 histone NTs other than H2BNT in H4acNuc (Fig. 2a). In all classes, EM density was detected only for BRPH, as in P300_H2B_·H4acNuc. We determined the complex structures in which HAT is directed toward H3NT (P300_H3-I_·H4acNuc; Supplementary Table 1) or H2ANT (P300_H2A_·H4acNuc). In all complexes, as in P300_H2B_·H4acNuc, bromodomain of P300_BRPH_ bound to H4K12ac of H4NTac, but the position of binding to the nucleosome was different for each (Fig. 2b). In P300_H2B_·H4acNuc and P300_H2A_·H4acNuc, bromodomain binding to H4NTac interacted with the minor groove at the superhelical location (SHL) +2. In P300H3-ioH4acNuc and P300_H3-II_·H4acNuc, it interacted with the minor groove at SHL +1. Importantly, in all structures except P300_H2B_·H4acNuc, HAT of P300_BRPH_ interacted with DNA through one or two basic patches (Fig. 2c). Of these, one basic patch^23^ (KJ basic patch; Supplementary Fig. 6) always interacted with the nucleosomal DNA. Along with that, another basic patch (KN basic patch) interacted with the linker DNA in P300_H3-I_·H4acNuc, suggesting an acetylation mechanism for H3NT that is less dependent on pre-acetylation of H4NT. This interaction mechanism explains why H3NT closest to the linker DNA is more likely to be acetylated indiscriminately.

**Fig. 2.**
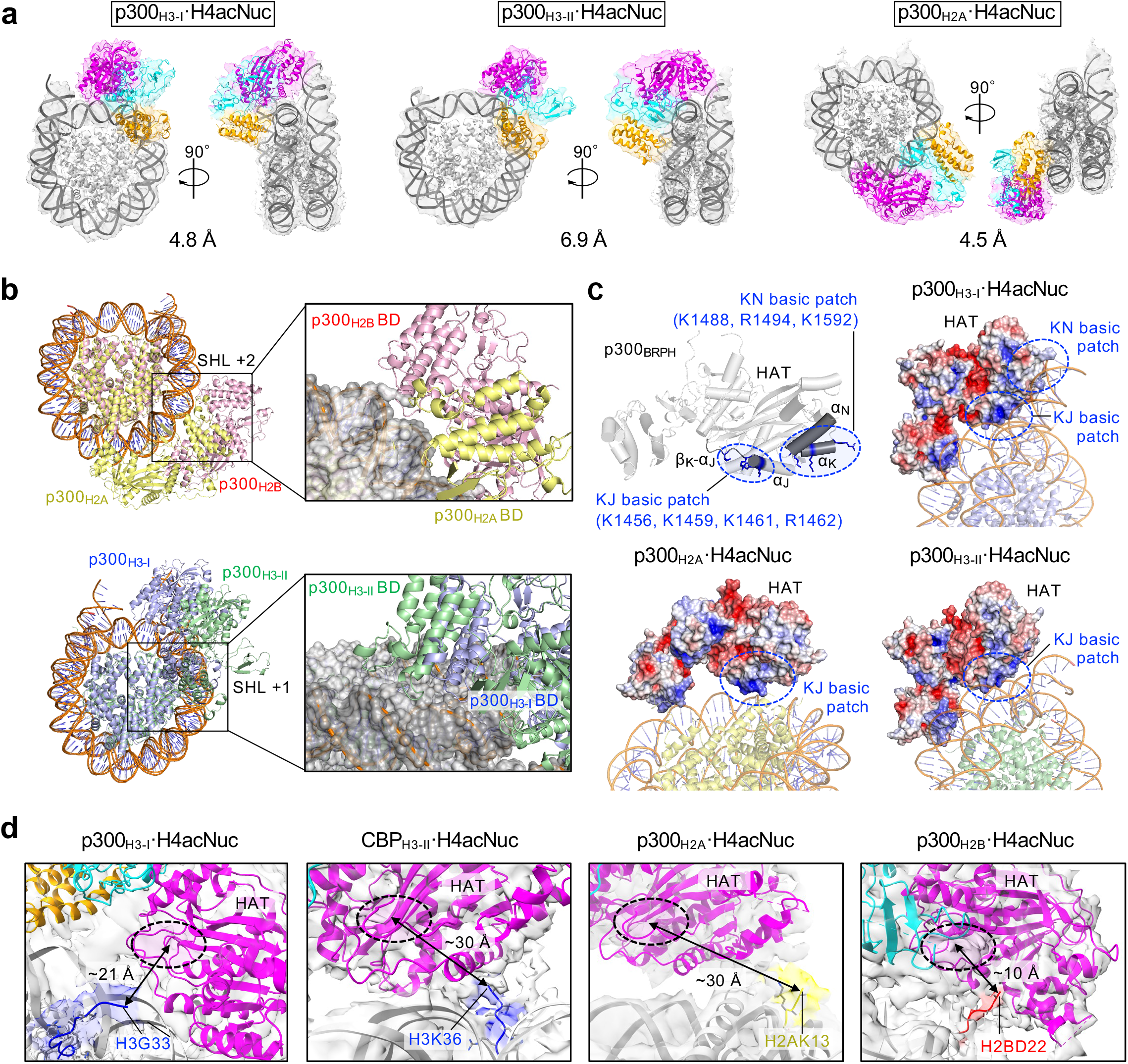
Modes of binding to multiple histone N-terminal tails. **a** Various conformations of p300_BRPH_ with the H4-di-acetylated nucleosome (H4acNuc) shown by cryo-electron microscopy (cryo-EM) maps and structural modelling: left, P300_H3-I_ (#5 in Supplementary Fig. 8); center, P300_H3-II_ (#6); right, P300_H2A_ (#7). (See Fig. 1 for color coding). **b** The positions of the superhelical location (SHL) at which p300 bromodomain (BD) interacts. Complex structures showing (top) superimposition of p300_H2B_ -H4acNuc (#4) and P300_H2A_-H4acNuc (#7) and (bottom) superimposition of p300_H3-I_-H4acNuc (#5) and p300_H3-II_-H4acNuc (#6). The respective regions where p300 BD interacts with DNA are indicated by black squares and are shown on the right in close-up, displaying p300 (ribbon diagram) and nucleosome (surface diagram). **c** Basic patches interacting with DNA at p300 HAT. In the top left panel (#5), two basic patches are circled in blue. One basic patch (K1456, K1459, K1461, and R1462) is located around the β_κ_-α_j_ loop (KJ basic patch) and the other (K1488, R1494, and K1592) is located in α_K_ and α_N_ (KN basic patch). The K/R residues involved in the interaction with DNA are shown in blue. The other three panels show the surface electrostatic potential of p300_BRPH_ for each complex structure, with surfaces charged positively in blue or negatively in red. Other panels (#5–#7): surface electrostatic potential of p300_BRPH_ for each complex structure. Positively charged surfaces are colored in blue and negatively charged surfaces in red. **d** Close-up views of the density and model structure of each NT in the H4acNuc complex. From left to right, the HAT catalytic center of p300 or CBP is shown in close proximity to H3NT (H3-I, #2), H3NT (H3-II, #10), H2ANT (#7), and H2BNT (#4) in H4acNuc. The rightmost panel showing H2BNT is another angle of Fig. 1h. Color codes of NT: blue: H3NT, yellow: H2ANT, red: H2BNT; cyan, p300 RP; magenta, p300 HAT.

In P300_H3-I_·H4acNuc, the density of H3NT toward HAT could be modeled for the C-terminal residues after G33 (Fig. 2d, leftmost panel). Since the distance between the substrate-binding pocket of HAT and G33 is ~21 Å, P300_BRPH_ of this complex is assumed to acetylate from the N-terminal side of H3 up to K23. In P300_H2A_·H4acNuc, the density of H2ANT toward HAT could be modeled for the C-terminal residues after K13 (Fig. 2d, second panel from the right). The distance between the substrate-binding pocket and K13 (~3Ü Å) suggests that P300_BRPH_ acetylates only K5 of H2A.

By preparing catalytically-active CBP_BRPHZT_ (residues 1084–1873) protein, we also determined three complex structures (Supplementary Fig. 1e, 7, 8, Supplementary Table 1) in which HAT of CBP_BRPH_ is oriented toward H2BNT (CBP_H2B_·H4acNuc) or H3NT (CBP_H3-I_·H4acNuc and CBP_H3-II_·H4acNuc). The overall conformation of CBP_BRPH_·H4acNuc complexes was almost identical to that of P300_BRPH_·H4acNuc, and the H4acNuc-binding modes of bromodomain and HAT were also almost identical to those of the corresponding P300_BRPH_·H4acNuc structures (Supplementary Fig. 9). Whereas the inter-molecular arrangement of P300_H3-II_·H4acNuc structure was determined only at 6.9 Å, the CBP_H3-II_·H4acNuc structure was determined at 4.2 Å, so the domain arrangement of CBP_BRPH_ and the structure of the H4acNuc-interactive region in its bromodomain could be determined. HAT of CBP_BRPH_ in CBP_H3-II_·H4acNuc was located closer to H3NT than in CBP_H3-I_·H4acNuc. The density of H3NT toward HAT could be modeled for the C-terminal residues after K36 (Fig. 2d, second panel from the left). The distance between the substrate-binding pocket and K36 (~30 Å) suggests that CBP_BRPH_ of this complex acetylates K27 in addition to lysine residues up to K23. Collectively, these results suggest that not only p300 but also CBP can acetylate multiple non-H4 histone NTs by rotating themselves on the nucleosome with their bromodomain bound to H4NTac as the axis.

### Bromodomain-dependent rotation of p300/CBP

Next, we examined why p300_BRPH_ or CBP_BRPH_ structures can direct HAT to such a variety of histone NTs when bound to H4acNuc. In all the structures obtained (five structures of p300_BRPH_·H4acNuc and three structures of CBP_BRPH_·H4acNuc), their bromodomain bound to H4K12ac of H4acNuc at the inside of its pocket and to the minor groove of the nucleosomal double-stranded DNA at the outside of the pocket (Fig. 3a). The position of bromodomain interacting with the minor groove varied from complex to complex, but in all structures, the third basic patch present around the BC-loop (BC basic patch; Fig. 3a, Supplementary Fig. 6) interacted electrostatically with the phosphate groups of the double-stranded DNA backbone. The electrostatic interaction both stabilizes binding to H4NTac *via* the inside of the bromodomain pocket and alters the relative positioning of HAT within the nucleosome, depending on where it occurs. Interestingly, neither p300_BRPH_ nor CBP_BRPH_ recognized any DNA sequences. Consequently, having DNA sequence–independent multivalent modes of binding to the nucleosome presumably allows HAT of p300/CBP to successively acetylate any of the non-H4 histone NTs within the H4NT-acetylated nucleosome.

**Fig. 3.**
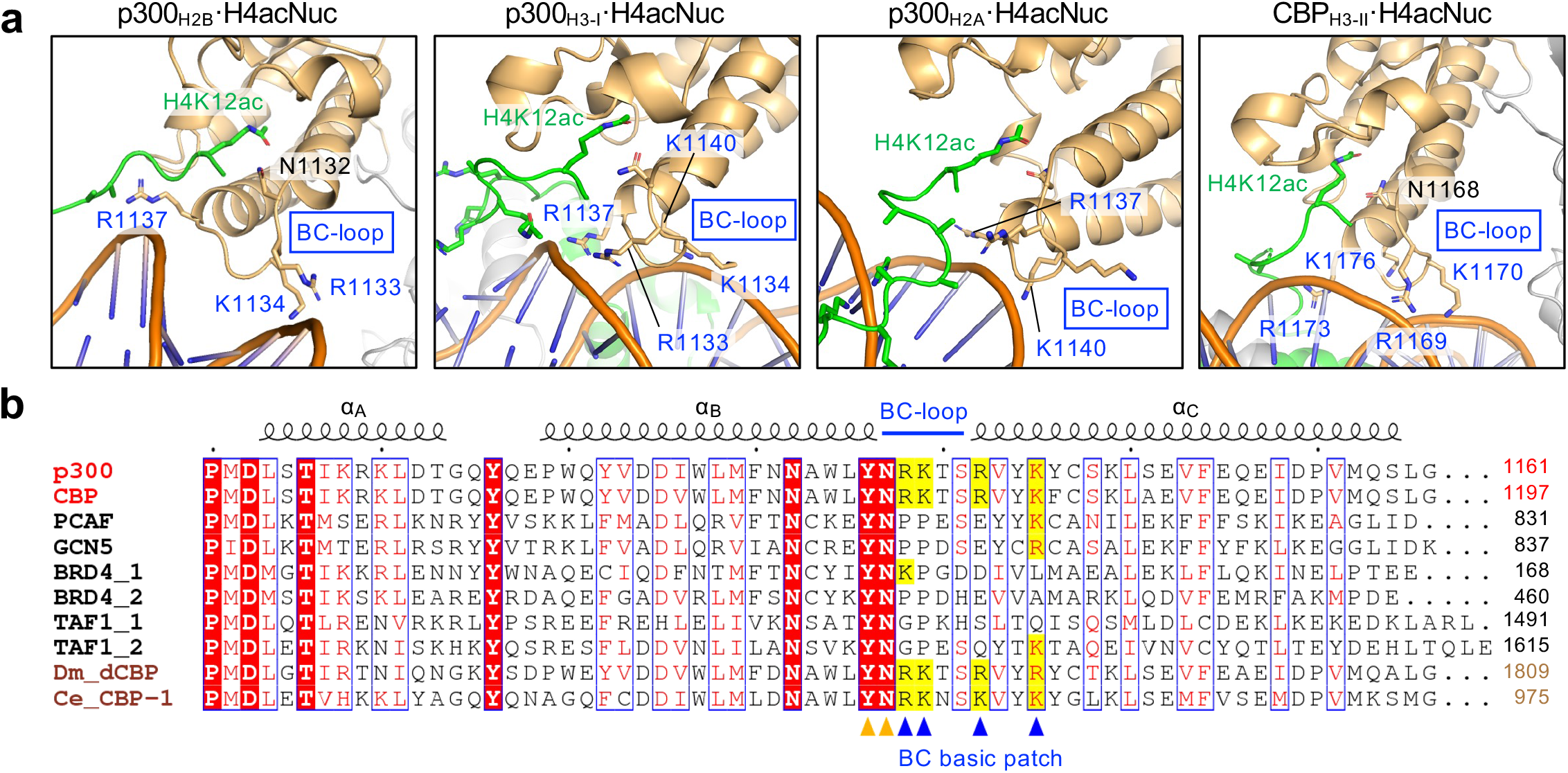
Rotational positions of p300_BRPH_/CBP_BRPH_ on the nucleosome fixed by a bromodomain loop. **a** Basic patch interacting with DNA at the p300/CBP bromodomain. Close-up views (#1, #5, #7, and #10 in Supplementary Fig. 8) of the bromodomain interacting with the H4K12acK16ac region of H4-di-acetylated nucleosome (H4acNuc) in complex with p300_BRPH_ are shown. This basic patch (R1133, K1134, R1137, and K1140 of p300; R1169, K1170, R1173, and K1176 of CBP) is located around the BC-loop of the p300/CBP bromodomain (BC basic patch; indicated by blue arrowheads at the bottom). The K/R residues involved in the interaction with DNA are shown in blue. In all structures, multiple R/K residues are involved in the interaction with DNA. Color code: light orange, the p300/CBP bromodomain; green, K12/K16-acetylated H4. **b** Sequence alignment around the BC-loop of the p300/CBP bromodomain. The positions of the BC-loop and three α-helices composing the bromodomain are shown on the top in blue and black, respectively. Protein names of representative human bromodomains are shown in black on the left. The protein names in brown are *D. melanogaster* Nejire/dCBP and *C. elegans* CBP-1. Residue numbers on the C-terminal side are shown on the right. Conserved or similar residues are shown in red and surrounded by blue boxes. The completely conserved residues are shown in white letters on a red background. The positions of residues involved in the recognition of acetyllysine inside the bromodomain pocket (Y1131 and N1132 of human p300) are indicated by orange arrowheads at the bottom. The positively charged residues conserved in the BC basic patch are indicated by a yellow background.

Sequence alignment of bromodomains indicates that R/K residues composing the BC basic patch (*i.e*., RKxxRxxK in p300/CBP, where x indicates an unrelated residue) are conserved only in p300 and CBP among all 61 human bromodomains (Fig. 3b, Supplementary Fig. 10). Importantly, these four R/K residues are conserved among metazoan p300 homologs, including *D. melanogaster* Nejire/dCBP^26^ and *C. elegans* CBP-1^27^.

### The BC basic patch is critical for ‘read/write’

To investigate the function of the BC basic patch in the ‘read/write’ mechanism, we prepared mutant proteins of p300_BRP_ or p300_BRPHZT_ in which all four positively charged residues were mutated with alanine (4A) or glutamic acid (4E) residues, respectively. First, concerning the ‘read’ mechanism, we measured the half-saturation concentration (*K*_1/2_) of p300_BRP_ for nucleosome binding by microscale thermophoresis with and without H4NTac (*i.e*., K12ac/K16ac), p300 4A mutation, and CBP30 (Supplementary Table 2). Wild-type p300_BRP_ bound to the unmodified nucleosome relatively strongly with a *K*_1/2_ of 2.2 nM. As expected, this binding was enhanced 6-fold by the presence of H4NTac (*K*_1/2_ = 0.35 nM), which was largely canceled by prior incubation with CBP30 (*K*_1/2_ = 1.2 nM). By contrast, 4A weakened the affinity for H4acNuc by 5-fold compared to the wild-type (*K*_1/2_ = 1.8 vs. 0.35 nM). Pre-incubation with CBP30 further weakened this affinity by 2-fold (*K*_1/2_ = 3.5 nM). These results reinforce our structures, in which the binding of p300_BRPH_ to H4acNuc is mediated both inside and outside the bromodomain pocket.

We next examined the effects of the two mutations of p300_BRPHZT_ on the H4NTac-dependent p300 ‘read/write’ mechanism (Fig. 4a, Supplementary Fig. 11). Both mutations dramatically reduced the H4NTac-dependent acetylation of H2BNT at all residues examined. They also reduced the H4NTac-dependent H3NTac found at K14, K23, and K27. This reduction was more prominent for 4E mutation, in which positively charged residues became negatively charged, than for 4A, in which residues became uncharged. H2BNTac was markedly reduced by CBP30 alone, and H3NTac was also almost completely suppressed by combining it with either mutation. These results suggest that the multivalent binding of p300_BRPHZT_ to H4acNuc is critical for Kac propagation to H2BNT and H3NT. Interestingly, both mutations also markedly reduced the H4NTac-independent acetylation by p300_BRPHZT_ (*i.e*., H2AK5ac, H3K18ac, and H4K5ac). Collectively, the BC basic patch is suggested to help p300 bind to and acetylate the nucleosome not only in an H4NTac-dependent manner but also independently of histone NTac.

**Fig. 4.**
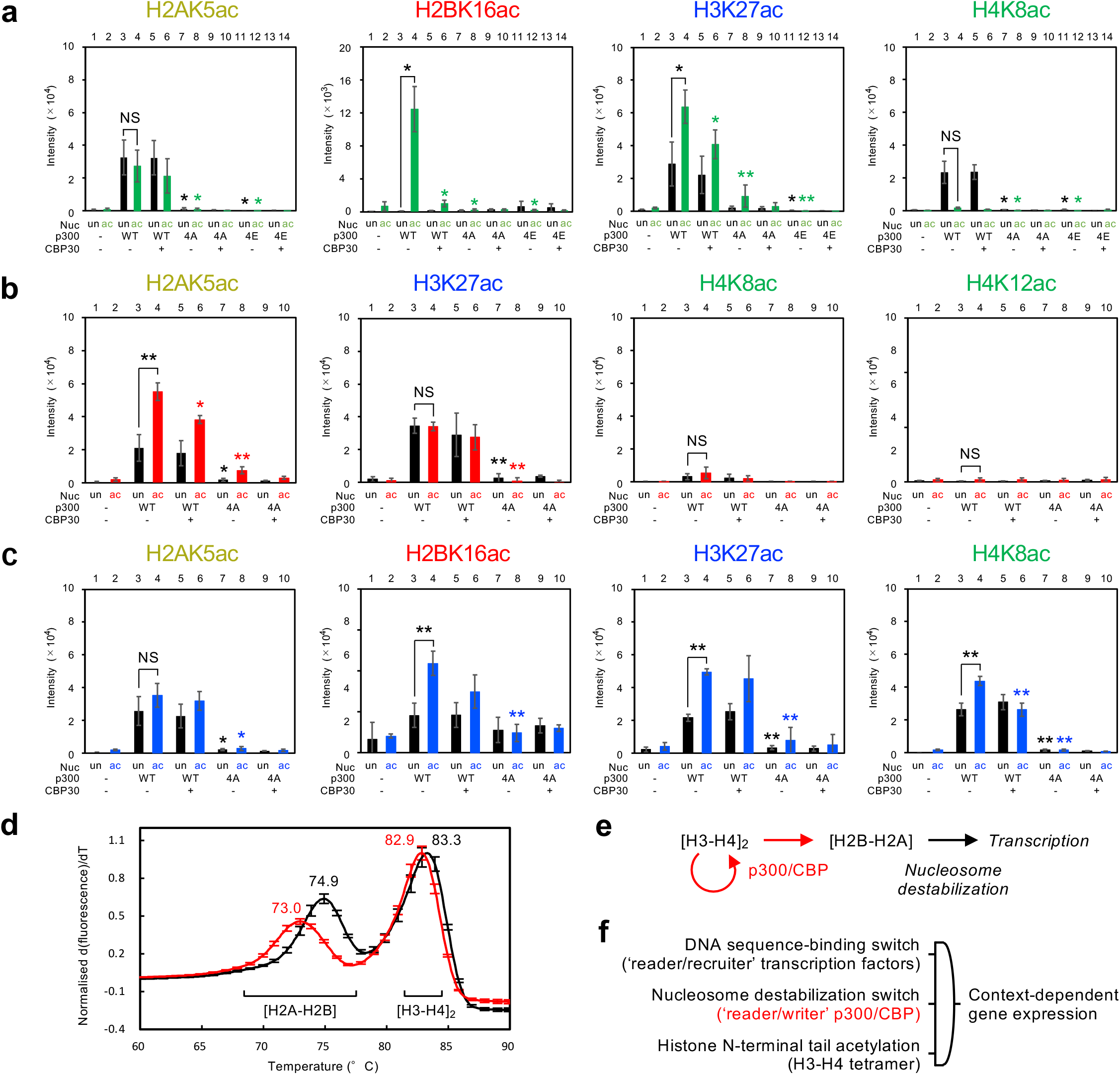
Propagation of intranucleosomal histone acetylation by p300_BRPHZT_. **a** *In vitro* acetyltransferase activity of p300_BRPHZT_ toward the H4-di-acetylated nucleosome. The histone and its residue at which acetylation was detected by immunoblotting are shown above each panel. Nucleosome (Nuc): un, unmodified; ac (green), H4K12/K16-acetylated. p300_BRPHZT_ (p300): WT, wild-type; 4A, with mutations of R1133A, K1134A, R1137A, and K1140A; 4E, with mutations of R1133E, K1134E, R1137E, and K1140E. CBP30: -, none; +, 10 μM. The y-axis indicates the immunoblotting signal intensity at 1 min after the reaction. Means ± SD (*N* = 3). Statistical significance was assessed by a two-sample one-sided Welch’s *t*-test (NS, *P* ≥ 0.05; \**P* < 0.05; \*\**P* < 0.01). The alternative hypothesis is as follows: lane 4, increase vs. lane 3; lanes 5, 7, and 11, decrease vs. lane 3; lanes 6, 8, and 12, decrease vs. lane 4). **b** *In vitro* acetyltransferase activity of p300_BRPHZT_ toward the H2B-tetra-acetylated nucleosome. Columns marked ac (in red) indicate the H2BK12/K15/K20/K23-acetylated nucleosome. Other indications are the same as in **a**. **c** *In vitro* acetyltransferase activity of p300_BRPHZT_ toward the H3-di-acetylated nucleosome. Columns marked ac (blue) indicate the H3K14/K18-acetylated nucleosome. **d** Thermal stability assay of the H2B-acetylated nucleosome. Mean values of thermal denaturation curves from 60.0 °C to 90.0 °C for derivative fluorescence intensity are plotted for the unmodified nucleosome (black line) and the H2BK12/K15/K20/K23-acetylated nucleosome (red line). The temperature at which the H2A-H2B dimer or the H3-H4 tetramer dissociates from the nucleosome is shown at the bottom. Means ± SD (*N* = 3). **e** ‘Epi-central’ model of histone acetylation signalling. Arrows indicate the flow of information, with acetylation information in red. **f** Hypothetical logic of context-dependent gene expression in metazoans. The symbol in the center indicates a triple-input AND logic gate.

### Directionality of histone NT acetylation

When acetylation was present in H4NT, p300_BRPHZT_ facilitated multisite lysine acetylation of the NTs in H2B and H3 within the nucleosome in a bromodomain-dependent manner. So, when acetylation is present in the non-H4 histone NTs, would p300 facilitate the acetylation of other histone NTs? To this end, we reconstituted the nucleosomes containing either H2B acetylated at K12/K15/K20/K23 (H2BacNuc) or H3 acetylated at K14/K18 (H3acNuc) and examined whether p300_BRPHZT_ acetylates them (Supplementary Fig. 12). H2BacNuc did not significantly facilitate p300_BRPHZT_-catalyzed NTac at the one-minute timepoint, except H2AK5ac (Fig. 4b, Supplementary Fig. 13). On the other hand, H3acNuc significantly facilitated p300_BRPHZT_-catalyzed H2BNTac and H4NTac at the one-minute timepoint (Fig. 4c, Supplementary Fig. 14, 15a). In all cases, including no pre-acetylation, p300_BRPHZT_ did not acetylate H4K16 at all and hardly acetylated H4K12. Collectively, the directionality in which p300_BRPHZT_ ‘reads/writes’ Kac between NTs falls into two types: bidirectional between H4NT and H3NT and unidirectional (from former to latter) between H4NT and H2BNT or between H3NT and H2BNT. This suggests that p300 bromodomain binds to H4NTac and H3NTac as reported^13^, but cannot or very weakly binds to H2BNTac.

Interestingly, when the nucleosome was pre-acetylated at H3K14/K18, p300_BRPHZT_ significantly propagated acetylation to K23/K27 (Fig. 4c, Supplementary Fig. 14). Since it is structurally difficult for p300/CBP to simultaneously ‘read/write’ K14/K18 and K23/K27 in one H3NT, this result suggests that p300_BRPHZT_ propagates lysine acetylation across the two H3NTs. Indeed, our model suggested that the H3NT pair in the nucleosome is close enough in proximity that p300_BRPH_ can ‘read/write’ H3NT(ac) on both (Supplementary Fig. 15b).

### H2BNTac destabilizes the nucleosome

When the multisite H2BNTac is triggered by H3NTac or H4NTac, are there any proteins recruited subsequently? The multisite H4NTac, such as H4K5ac/K8ac, is a scaffold to which the bromodomains of proteins in the bromodomain and extra-terminal (BET) family preferentially bind^28, 29^. Since the sequence between K5ac and K8ac of H4NT is similar to that between K12ac and K15ac of H2BNT^30^, the BET bromodomains may bind to the di-acetylated H2BNT. To this end, we examined dissociation constants (*K_D_*) between p300 or BET bromodomains and several di-acetylated H2BNT or H4NT peptides by isothermal titration calorimetry (Supplementary Table 3). BRD4_BD1_, the N-terminal bromodomain of BET family protein BRD4, bound very weakly or poorly to the di-acetylated H2B peptides tested, with a minimal *K_D_* of 430 μM (*i.e*., for K12ac/K15ac). This affinity was 20-fold weaker than the *K_D_* between BRD4_BD1_ and the H4K12ac/K16ac peptide (22 μM). Hence, it is unlikely that multisite H2BNTac could recruit BET proteins as the multisite H4NTac does. Additionally, bromodomain-containing p300_BRP_ also bound very weakly or poorly to the di-acetylated H2B peptides (K20ac/K23ac), showing a minimal *K_D_* of 200 μM. This affinity was also 13-fold weaker than the *K_D_* between p300_BRP_ and the H4K12ac/K16ac peptide (15 μM). Therefore, multisite H2BNTac is an unlikely scaffold for nucleosome binding either by p300 or BRD4 bromodomains. This is consistent with the fact that the multisite H2BNTac did not facilitate the p300_BRPHZT_-catalyzed histone NTac, except for H2AK5ac.

Thus, multisite H2BNTac may serve as an endpoint of this signaling rather than recruiting other proteins. If so, it is natural to assume that the high correlation between H2BNTac and the enhancer activity^16^ is not the result of transcription-coupled histone exchange, but rather its direct cause. Based on this hypothesis, we examined the effect of H2BNTac on the thermal stability of the nucleosome. Of the stepwise dissociation of the H2A-H2B dimer and the H3-H4 tetramer, the lower *T*_m_, reflecting the dissociation of the H2A-H2B dimer, decreased 0.6 °C for H3acNuc and increased 0.1 °C for H4acNuc compared to the unmodified nucleosome (Supplementary Fig. 16), but decreased 1.9 °C for H2BacNuc (73.0 vs. 74.9 °C; Fig. 4d). The higher *T*_m_, reflecting the dissociation of the H3-H4 tetramer, was similar to that in the unmodified nucleosome at any acetylation (82.9–83.1 °C vs. 83.3 °C for the unmodified nucleosome). These results suggest that multisite H2BNTac selectively promotes H2A-H2B dissociation from the nucleosome.

## Discussion

Since the study of Allfrey et al.^5^, histone acetylation has become recognized as an important PTM regulating eukaryotic gene transcription. The present study revealed how p300/CBP recognizes H4NTac and propagates lysine acetylation to non-H4 histones within the nucleosome. That is, p300/CBP ‘reads’ H4NTac at the bromodomain pocket, ‘rotates’ in multiple directions, and rapidly ‘writes’ NTac to non-H4 histones within the nucleosome independently of the DNA sequence. To our knowledge, this is the first structural evidence showing how a particular PTM in the nucleosome is ‘read/written’ by the enzyme to self-propagate. In contrast to the mechanisms by which histone methylation at H3K9 and H3K27 involved in transcriptional repression spreads to neighbouring nucleosomes^31^, histone acetylation involved in transcriptional activation spreads within a single nucleosome. The ‘read/write’ role of p300/CBP derived from our data is twofold: 1) ‘replication’ of NTac within the H3-H4 tetramer, and 2) ‘transcription’ of NTac from the H3-H4 tetramer to the H2B-H2A dimer.

The first role for p300/CBP was derived from the symmetry in the flow of acetylation information between H3NT and H4NT (Supplementary Fig. 15a). The distance between the H4NT pair within the nucleosome was too far for p300/CBP to ‘read/write’ Kac directly between them. However, when bound to one of a pair of H4NTs, p300/CBP was at the right proximity to acetylate the closer NT for each of the pairs of H3NT, H2BNT, and H2ANT. Our data also suggest that p300_BRPH_ can propagate Kac between the H3NT pair (Supplementary Fig. 14, 15b). Therefore, p300/CBP may ‘read/write’ the multisite NTac 1) from one of the H4NT pair to the proximal H3NT, 2) between the H3NT pair, and 3) from the distal H3NT to the other H4NT. p300/CBP would be a maintenance acetyltransferase that would ensure self-perpetuation of Kac in the H3-H4 tetramer. Based on our data and previous reports^13, 15, 32, 33^, possible hypotheses include: 1) H3K14ac/K18ac self-perpetuates by p300/CBP as flexible epigenetic storage; 2) H4K8ac/K12ac self-perpetuates with the help of H3K14ac/K18ac as robust epigenetic storage; and 3) H3K14ac and H4K8ac catalyzed by p300/CBP are mitotic bookmarks supporting the self-perpetuation mechanisms.

The second role for p300/CBP was derived from the asymmetry in the flow of acetylation information from H4NT or H3NT to H2BNT. p300/CBP interacts with at least 400 proteins, including various DNA-binding transcription factors (TFs)^34^. Presumably, p300/CBP binds to the partner TFs bound to specific DNA sequences *via* its intramolecular domain(s) such as TAZ2, while simultaneously interacting *via* bromodomain and ZZ^35^ with nucleosomes not only proximal to the enhancers but also distal ones by chromatin looping. Then, p300/CBP acetylates H2B (*e.g*., at active enhancers, their target promoters, and gene body regions) of those nucleosomes when H3NTac or H4NTac is present. This would promote dissociation of the H2A-H2B dimer from the nucleosome in these regions, at which RNAPII enters DNA more readily and/or transcribes it more productively. In this light and given other findings on the behaviour of H2B^36, 37, 38, 39^, it makes sense that p300-catalyzed H2BNTac is the genuine signature of active enhancers and their target promoters^16^. Since RNAPII complexed with the histone chaperone FACT flips the histone octamer and exchanges one H2A-H2B dimer during traverse across the nucleosome^40^, the H2BNTac information ‘written’ by p300/CBP should be ‘erased’ by subsequent successive transcription.

The asymmetry in the flow of acetylation information would originate from a KK sequence unique to H2BNT (Supplementary Fig. 12b). Crystal structures^12, 13^ suggest that a residue having a long side chain such as lysine (K) just before or after Kac prevents p300 bromodomain binding, and indeed this sequence is absent in H3NT and H4NT. H2BNT is less conserved than H3NT and H4NT, but two KK sequences are conserved in H2B from yeast to human, a feature not found in other histones^16^. This KK sequence may be a mechanism devoted to nucleosome destabilization *via* dense multisite lysine acetylation, preventing unnecessary signaling *via* the Kac ‘readers’. Also, acetylation of H2BNT was structurally the least dependent on the DNA binding activity of p300 HAT among the histone NTs (Fig. 2b, c). This suggests that H2BNTac is the least indiscriminately acetylated, strictly p300 ‘reader’ activity-regulated ‘read/write’ switch with the best signal-to-noise ratio.

The specificity of chromatin acetylation has been attempted to be explained primarily by one of the following two models: 1) a TF binding to a specific DNA sequence specifies the chromatin sites where its partner acetyltransferase (complex) acetylates, or 2) an acetyltransferase (complex) binding to histone NTac specifies the chromatin sites to acetylate independently of DNA sequence. The current situation is more supportive for the former because the epigenomic locations of p300 largely overlap with those of its partner TFs independently of its acetyltransferase activity^20^, and the acetyltransferase activity of p300 directly depends on activation of TF ligands^13^. On the other hand, support for the latter is that p300 bromodomain is critical in modulating its enzymatic activity and its association with chromatin^35^ and also that inhibition of the p300/CBP bromodomain pockets decreases H3K27ac and transcription of enhancer-proximal genes^41^. Based on the present data, we support a model that integrates both. That is, the global specificity of chromatin acetylation would be determined by the proximity or contactable range of the chromatin to p300/CBP, which is recruited by its partner TF bound to a specific DNA sequence in the genomic DNA. Subsequently, the local specificity of chromatin acetylation would be determined by whether the H3-H4 tetramer is pre-acetylated (even slightly) more than usual when p300/CBP interacts with each nucleosome of that chromatin.

Finally, we propose a model of p300/CBP-catalyzed histone acetylation signaling (Fig. 4e). Here, acetylations of the H3-H4 tetramer and the H2A-H2B dimer play distinct roles. Just as DNA is the genetic storage, p300/CBP ‘replicates’ histone acetylation within the H3-H4 tetramer, which self-perpetuates as the epigenetic storage. Also, just as RNA is the genetic processor, p300/CBP ‘transcribes’ histone acetylation from the H3-H4 tetramer to the H2B-H2A dimers in a strictly regulated manner, which then self-sacrifices to express specific genes as the epigenetic processor. Regarding epigenetic switches, a DNA sequence–binding protein that may self-perpetuate by a positive feedback mechanism is a ‘read/recruit’ switch that specifies a gene to be transcribed from genomic DNA^42, 43^, and p300/CBP is presumably a nucleosome destabilization ‘read/write’ switch for productively transcribing that gene from the nucleosomes (Fig. 4f). The logic of context-dependent gene expression in metazoans would be a triple-input AND-gated circuit consisting of the DNA sequence–binding switch^42^, the nucleosome destabilization switch, and histone N-terminal tail acetylation of the H3-H4 tetramer^4, 5^.

## Methods

### Expression and purification of p300 and CBP

The cDNA sequences encoding human p300_BRPHZT_ (residues 1048–1836) and CBP_BRPHZT_ (residues 1084–1873) were amplified by PCR and subcloned into a pFastBac HT vector with a glutathione S-transferase (GST)- encoding sequence inserted after an N-terminal polyhistidine tag. Site-directed mutagenesis of the p300_BRPHZT_ 4A-substituted (R1133A, K1134A, R1137A, and K1140A) and 4E-substituted (R1133E, K1134E, R1137E, and K1140E) mutant proteins was performed by PCR, using the *DpnI* restriction enzyme. The p300_BRPHZT_, CBP_BRPHZT_, and p300_BRPHZT_ 4A- and 4E-substituted mutant proteins were expressed in baculovirus-infected High Five insect cells (Thermo Fisher Scientific). Baculoviruses were produced using the Bac-to-Bac baculovirus expression system (Invitrogen). The baculovirus-infected cells were collected 72 hrs after transfection, and cell pellets were frozen at −80 °C until purification. Frozen High Five cells were resuspended in 20 mM Tris-HCl buffer (pH 7.2) containing 500 mM NaCl, 10% glycerol, 20 mM imidazole, 0.1% NP-40, 1.5 mM MgCl_2_, 1 μM ZnCl_2_, DNase I (Sigma Aldrich), and cOmplete (EDTA-free) Protease Inhibitor Cocktail (Roche). Cells were lysed by sonication and clarified by centrifugation. The cell lysate from each sample was loaded onto a HisTrap HP column (GE Healthcare) and eluted using 50 mM Tris-HCl buffer (pH 8.0) containing 500 mM NaCl, 10% glycerol, and 500 mM imidazole. After buffer exchange using a HiTrap Desalting column (GE Healthcare), the N-terminal polyhistidine-GST tag was cleaved by incubation with TEV protease at 4 °C overnight. The cleaved protein was then reapplied to a HisTrap HP column, and the flow-through fraction was collected. The collected fractions were purified by size-exclusion column chromatography, using a HiLoad Superdex 200 26/60 (GE Healthcare) equilibrated with 20 mM HEPES buffer (pH 7.2) containing 250 mM NaCl, 1 mM Tris (2-carboxyethyl) phosphine (TCEP), and 5 μM ZnCl2. Purified protein was concentrated using an Amicon Ultra-15 centrifugal filter unit (Millipore, 50 kDa MWCO) and flash frozen in liquid nitrogen.

For microscale thermophoresis measurements and isothermal titration calorimetry, the cDNAs encoding p300_BRP_ (residues 1048–1282) were amplified by PCR and subcloned into the pET28a(+) vector encoding GST and the polyhistidine tag. Site-directed mutagenesis of p300_BRP_ 4A-substituted mutant was performed by PCR using the *DpnI* restriction enzyme. All introduced mutations were verified by DNA sequencing. The wild-type p300_BRP_ and 4A- substituted mutant were expressed in LB broth of *E. coli* BL21 (DE3) cells at 37 °C until the OD_600_ reached 0.8. The temperature was then shifted to 18 °C, and isopropyl-β-D-thiogalactopyranoside was added to a final concentration of 300 μM to induce protein expression. The cultures were incubated for an additional 20 hrs and collected by centrifugation. Cell pellets were resuspended in 50 mM Tris-HCl buffer (pH 8.0) containing 500 mM NaCl, 10% glycerol, 20 mM imidazole, and 10 μM ZnCl_2_. Cell lysates prepared by sonication and centrifugation were purified on a HisTrap HP column (GE Healthcare). The N-terminal polyhistidine-GST tag was cleaved by incubation with TEV protease at 4 °C overnight. The cleaved protein was then reapplied to a GSTrap HP column (GE Healthcare), and the flow-through fraction was collected. The eluted fractions were loaded on a HiLoad Superdex 200 16/60 column (GE Healthcare) equilibrated with HEPES buffer (pH 7.4) containing 150 mM NaCl and 1 mM TCEP.

### Reconstitution of residue-specific acetylated nucleosomes

The recombinant human histone proteins containing residue-specific Kac(s) (K12/K15/K20/K23-acetylated H2B; K14/K18-acetylated H3; K12/K16-acetylated H4) were synthesized by genetic code reprogramming, essentially as described^44, 45^. Briefly, human H2B type 1-J, H3.1, or H4 cDNA with codons for the specified residues replaced with the TAG triplets and a terminal TAA stop codon was used for protein synthesis in the coupled transcription–translation cell-free system. The recombinant human unmodified histones H2A type 1-B/E, H2B type 1-J, H3.1, and H4 were expressed in *E. coli* and the synthesized histones with or without designed Kac(s) were purified as reported^44, 45^. Reconstituted nucleosomes consisted of the histone octamer with the designed Kac(s) and the palindromic 180-bp DNA that consisted of the 146-bp human α-satellite DNA and 17-bp linker DNA (5’-ATC CGT CCG TTA CCG CC-3’) linked at both ends^46^, and reconstitution was performed as reported. For cryo-EM analysis, reconstituted nucleosomes were dialyzed against HEPES buffer (pH 7.2) containing 150 mM NaCl.

### Immunoblot analysis

The histones containing residue-specific acetylations or the nucleosomes used for the acetyltransferase activity assay were electrophoresed in a 10–20% *sodium* dodecyl sulfate polyacrylamide gel (SDS-PAGE) (DRC, NXV-396HP20) and transferred onto a nitrocellulose membrane (BIO-RAD, 1620112) at 20 V for 10 min by the semi-dry method. The membrane was blocked with Bullet Blocking One for Western Blotting (Nacalai Tesque, 13779-01) for 10–30 min at 25 °C. Membranes were incubated for 20–40 min at 25 °C with Bullet ImmunoReaction Buffer (Nacalai Tesque, 18439-85) containing the following antibodies at the indicated dilution rate: H2AK5ac (Abcam, ab45152, 1/3,000), H2B (Cell Signaling, #12364, 1/1,000), H2BK12ac (Abcam, ab40883, 1/500), H2BK15ac (Abcam, ab62335), H2BK16ac (Abcam, ab177427, 1/1,000), H2BK20ac (Abcam, ab177430, 1/500), H2BK23ac (Abcam, ab222770, 1/1,000), the C-terminus of H3 (Merck, 07-690, 1/3,000), H3K14ac (Merck, 07-353, 1/1,000), H3K18ac (Abcam, ab1191, 1/1,000), H3K23ac (Merck, 07-355, 1/1,000), H3K27ac (Merck, 07-360, 1/3,000), the C-terminus of H4 (Abcam, ab10158, 1/1,000), H4K5ac (MABI, 0405, 1/500), H4K8ac (MABI, 0408, 1/500), H4K12ac (MABI, 0412, 1/500), or H4K16ac (MABI, 0416, 1/500). The membranes were then washed with TBS-T (4 times, for 5 min each time) and incubated with Bullet ImmunoReaction Buffer (Nacalai Tesque, 18439-85) containing peroxidase-conjugated anti-mouse IgG (GE Healthcare, NA931) or anti-rabbit IgG (GE Healthcare, NA934) for 20 min at 25 °C. The membranes were then washed with TBS-T (4 times, for 5 min each time) and were detected using enhanced chemiluminescence (Chemi-Lumi One Super: Nacalai Tesque, 02230-30). The immunoblotted membranes were imaged using ImageQuant LAS-4000 (GE Healthcare). From the images obtained, the immunoblotting signal intensity of each band was quantified using ImageJ (RSB, https://imagej.nih.gov), and the background intensity was subtracted.

### Acetyltransferase assay

Acetyltransferase activity assays of p300_BRPHZT_ against the nucleosome were performed at a molar ratio of p300_BRPHZT_:180-bp nucleosome:acetyl-CoA (Nacalai Tesque, 00546-54) = 1:1:10 in 10 μl of reaction solution containing 5.6 mM HEPES-NaOH (pH 7.4), 4.4 mM Tris-HCl (pH 7.4), and 130 mM NaCl. Briefly, in a 0.2-ml PCR tube (Thermo Fisher Scientific, 3414JP), 2.4 μl of 10 mM HEPES-NaOH (pH 7.4) buffer containing 10 pmol p300_BRPHZT_ and 300 mM NaCl was mixed with 3.2 μl of 10 mM HEPES-NaOH (pH 7.4) buffer containing 10 pmol nucleosomes (with or without the indicated residue-specific histone acetylations) and 150 mM NaCl, and 4.4 μl of 10 mM Tris-HCl (pH 7.4) buffer containing 100 pmol acetyl-CoA. Every minute for 0–3 min after the reaction at 37 °C, each sample was mixed with 20 μl of 4× SDS-PAGE loading buffer and immediately heated at 95 °C for 3 min to stop the reaction, and 30 μl of water was added. For each sample solution, 4 μl was electrophoresed in a 10–20% SDS Tris-Tricine polyacrylamide gel and applied for the immunoblot analysis. All assays were performed in triplicate.

### Cryo-EM sample and grid preparation

To prepare p300_BRPHZT_·H4acNuc and CBP_BRPHZT_·H4acNuc complexes, p300_BRPHZT_ (or CBP_BRPHZT_), acetyl-CoA, and H4acNuc were mixed at a molar ratio of 4:6:1 in 20 mM HEPES buffer (pH 7.2) containing 150 mM NaCl, 1 μM ZnCl_2_, and 1 mM TCEP and incubated for 30 min. To purify and crosslink the complex, the reaction mixture was fractionated by the GraFix method^47^. A gradient for GraFix was formed with a top solution of 10 mM HEPES buffer (pH 7.2) containing 50 mM NaCl, 1 mM TCEP, 10% glycerol, and 0.01% glutaraldehyde) and a bottom solution of 10 mM HEPES (pH 7.2) containing 50 mM NaCl, 1 mM TCEP, 30% glycerol, and 0.15% glutaraldehyde, using a Gradient Master (BioComp). The reaction mixture was applied onto the top of the gradient solution and was centrifuged at 41,000 rpm at 4 °C for 17 hrs using an SW41 Ti rotor (Beckman Coulter). The gradient was fractionated using a Piston Gradient Fractionator (BioComp). The fractions were examined by electrophoresis at 150 V for 50 min on 6% native TBE polyacrylamide gels, and then the fractions corresponding to H4acNuc complex were pooled. Glutaraldehyde was quenched by the addition of Tris-HCl at pH 7.6 to a final concentration of 100 mM. The samples were dialyzed against 10 mM HEPES buffer (pH 7.2) containing 50 mM NaCl and 1 mM TCEP and concentrated using Amicon Ultra 100K (Merck Millipore) for electron microscopy analyses. A 3.0-μL aliquot of the samples of the p300_BRPHZT_·H4acNuc complex (2.0 mg ml^-1^) or the CBP_BRPHZT_·H4acNuc complex (2.0 mg ml^-1^) was each applied to glow-discharged, holey, copper grids (Quantifoil Cu R1.2/1.3, 300 mesh) with a thin carbon-supported film. The grids were plunge-frozen into liquid ethane using Vitrobot Mark IV (Thermo Fisher Scientific). Parameters for plunge-freezing were set as follows: blotting time, 3 sec; waiting time, 3 sec; blotting force, −5; humidity, 100%; and chamber temperature, 4 °C.

### Cryo-EM data collection and image processing

Cryo-EM data were collected with a Tecnai Arctica transmission electron microscope (Thermo Fisher Scientific) operated at 200 kV using a K2 summit direct electron detector (Gatan) at a nominal magnification of 23,500× in electron-counting mode, corresponding to a pixel size of 1.47 Å per pixel. The movie stacks were acquired with a defocus range of −0.9 to −1.7 μm with total exposure time of 12 s fragmented into 40 frames with the dose rate of 50.0 e^-^/Å^2^. Automated data acquisition was carried out using SerialEM software^48^. The cryo-EM data were also collected with a Krios G4 transmission electron microscope (Thermo Fisher Scientific) operated at 300 kV using a K3 direct electron detector (Gatan) at a nominal magnification of 105,000×in electron-counting mode, corresponding to a pixel size of 0.83 Å) per pixel. The movie stacks were acquired with a defocus range of −0.8 to −2.0 μm with total exposure time of 2.3 s fragmented into 50 frames with the dose rate of 50.0 e^-^/Å^2^. These data were automatically acquired by the image-shift method using the EPU software. All cryo-EM experiments were performed at the RIKEN Yokohama cryo-EM facility. All image processing was performed with RELION-3.1^49^. Dose-fractionated image stacks were subjected to beam-induced motion correction using MotionCor2^50^ and the CTF parameters were estimated with CTFFIND-4.1^51^. Particles were automatically picked using crYOLO^52^ with a box size of 135 × 135 pixels for the Tecnai Arctica dataset and a box size of 225 × 225 for the Krios G4 dataset. These particles were extracted and subjected to several rounds of 2D and 3D classifications using RELION 3.1. The selected particles were then re-extracted and subjected to 3D refinement, Bayesian polishing^53^, and subsequent postprocessing of the map improved its global resolution, according to the Fourier shell correlation with the 0.143 criterion^54^. Details of the data collection and image processing are summarized in Supplementary Table 1 and Supplementary Fig. 4, 5, and 7.

### Model building and refinement

For p300_BRPH_, each domain of the p300_BRPH_ crystal structure (PDB ID: 6GYR) was divided and fitted to a cryo-EM map as a rigid body using “fit in map” in the visualisation software UCSF Chimera^55^. For CBP_BRPH_, as with p300, each domain of the crystal structure (BD and HAT; PDB ID: 5U7G) and AlphaFold structure (RP; AF-Q92793-F1) was fitted to the cryo-EM map. For H4acNuc, the crystal structure of the 146 bp nucleosome (PDB ID: 1KX3) was fitted to a cryo-EM map, and then linker DNA and H4NTac were manually modeled and each histone was substituted for an amino acid residue using Coot^56^. For all structures, when rigid bodies could not be fitted to the cryo-EM map, we performed a flexible fitting by using the plug-in ISOLDE^57^. The final model was refined by PHENIX^58^, and the stereochemistry was assessed by MolProbity^59^. Statistics for cryo-EM model refinement are summarized in Supplementary Table 1. All figures were generated using either UCSF Chimera (v1.15), UCSF ChimeraX (v1.4)^60^, or PyMOL (v2.5)^61^. The mapping of electrostatic potential was achieved using PyMOL with the Adaptive Poisson-Boltzmann Solver (APBS) Electrostatics plugin (https://pymolwiki.org/index.php/APBS_Electrostatics_Plugin).

### Microscale thermophoresis

Microscale thermophoresis was performed essentially as previously described^45^. Briefly, the wild-type p300_BRP_ and 4A-substituted mutant were each fluorescently labeled, using a His-tag Labeling Kit (NanoTemper Technologies, cat. no. MO-L018). For measurements using CBP30, CBP30 was added beforehand to the wild-type and the 4A-substituted mutant to a final concentration of 10 μM and incubated for at least 20 min. Labeled proteins and the nucleosomes were buffered with 10 mM HEPES-NaOH buffer (pH 7.4) containing 150 mM NaCl. Measurements were taken by using a Monolith NT.115 Instrument (NanoTemper Technologies) at 25 °C according to the manufacturer’s protocol. Each assay was performed in biological triplicate using the unmodified and H4NT-acetylated nucleosome. The measured data were fitted to the Hill equation using NT analysis software (NanoTemper Technologies).

### Isothermal titration calorimetry

The N-terminal bromodomain of human BRD4 (residues 44–168; BRD4_BD1_) and the BRP domain of p300 (residues 1048–1282; p300_BRP_) were purified essentially as described^45, 62^. H2B (residues 1–27) and H4 (1–20) peptides with indicated Kac(s) were purchased from Toray Research Center. Measurements were conducted at 25 °C in 10 mM HEPES-NaOH buffer (pH 7.4) containing 150 mM NaCl on a MicroCal Auto-iTC_200_ microcalorimeter (Malvern). Approximately 400 μl of 100 μM protein solution was loaded into the sample cell, and each of 1 mM acetylated histone NT peptides was loaded into an injection syringe. The collected data were analyzed using Origin 7 SR4, ver. 7.0552 software (OriginLab Corporation) supplied with the instrument to calculate enthalpies of binding (ΔH) and *K_D_*. A single binding site model was used in all assays.

### Nucleosome thermostability assay

The thermal stability of the nucleosome with specific residues acetylated and of the unmodified nucleosome was measured essentially as previously described^44^. Briefly, 20 pmol nucleosomes in 20 mM Tris-HCl buffer (pH 7.5) containing 1 mM EDTA and 1 mM dithiothreitol were reacted with a 4-fold concentration of Protein Thermal Shift Dye (Life Technologies) in a 20 μl reaction volume. Fluorescence intensity produced by the reaction was monitored (*λ*ex = 580 nm; *λ*em = 623 nm) every second from 25.0 °C to 99.9 °C with a temperature change of 0.9 °C/min, using a

QuantStudio 6 PCR system (Life Technologies). Data were normalized with the maximal fluorescence intensity as 100%.

### Multiple sequence alignment

The multiple sequence alignment was performed by Clustal W^63^ in UniProt (http://www.uniprot.org/align/) and was rendered by ESPript^64^ (http://espript.ibcp.fr/ESPript/ESPript/). Amino acid sequences used for the alignment are: human p300 (UniProt ID: Q09472), human CBP (Q92793), human PCAF (Q92831), human GCN5 (Q92830), human BRD4 (O60885), human TAF1 (P21675), *D. melanogaster* Nejire/dCBP (Q9W321), *C. elegans* CBP-1 (P34545), and *A. thaliana* PCAT2 (Q9C5X9). Amino acid sequences of other human bromodomain-containing proteins in Supplementary Fig. 10 were obtained from UniProt.

## Data availability

The cryo-EM maps and the corresponding atomic coordinates have been deposited in the Electron Microscopy Data Bank (https://www.ebi.ac.uk/pdbe/emdb/) and the Protein Data Bank (http://www.rcsb.org) under the accession codes shown in Supplementary Table 1.

## Acknowledgements

We thank the RIKEN Yokohama cryo-EM facility of the Center for Biosystems Dynamics Research (BDR) for their support in the cryo-EM data collection; Kazuharu Hanada, Mio Inoue, and Sayako Miyamoto-Kohno for sample preparation; Nando Dulal Das, Hideaki Niwa, Shinsuke Ito, and Haruhiko Koseki for discussions; Yuki Saito for clerical assistance; and Masami Horikoshi, Shigeyuki Yokoyama, and Minoru Yoshida for encouragement. This work was supported by Grants-in-Aid from the Japan Society for the Promotion of Science (JP19K16062 and JP21K15035 to M.K.; JP16H05089, JP20H03388, JP20K21406, and JP21H05764 to T.U.); the PRESTO program of the Japan Science and Technology Agency (JPMJPR12A3 to T.U.); the Platform Project for Supporting Drug Discovery and Life Science Research (BINDS) from Japan Agency for Medical Research and Development (JP21am0101082 to M.S.; JP21am0101115: support No. 2959); the ‘Structural Cell Biology Project’ of RIKEN BDR to M.K.; the ‘Epigenome Manipulation Project’ of the All-RIKEN Projects to T.U.; and the RIKEN Pioneering Project ‘Genome Building from TADs’ to T.U.

## Author contributions

M.K. designed the research, performed the cryo-EM structure and biochemical analyses, and drafted the structural part of the manuscript. S.M., M.W., and S.S. performed the biochemical analysis. T.U-K. assisted the cryo-EM measurement. M.S. assisted the protein preparation. T.U. conceived and supervised the project, designed the research, and wrote the manuscript.

## Additional information

**Supplementary information** is available for this paper.

**Correspondence** and requests for materials should be addressed to Takashi Umehara.

