## Supplementary Information for "Epigenetic mechanisms to propagate histone acetylation by p300/CBP"

### Supplementary Figure 1

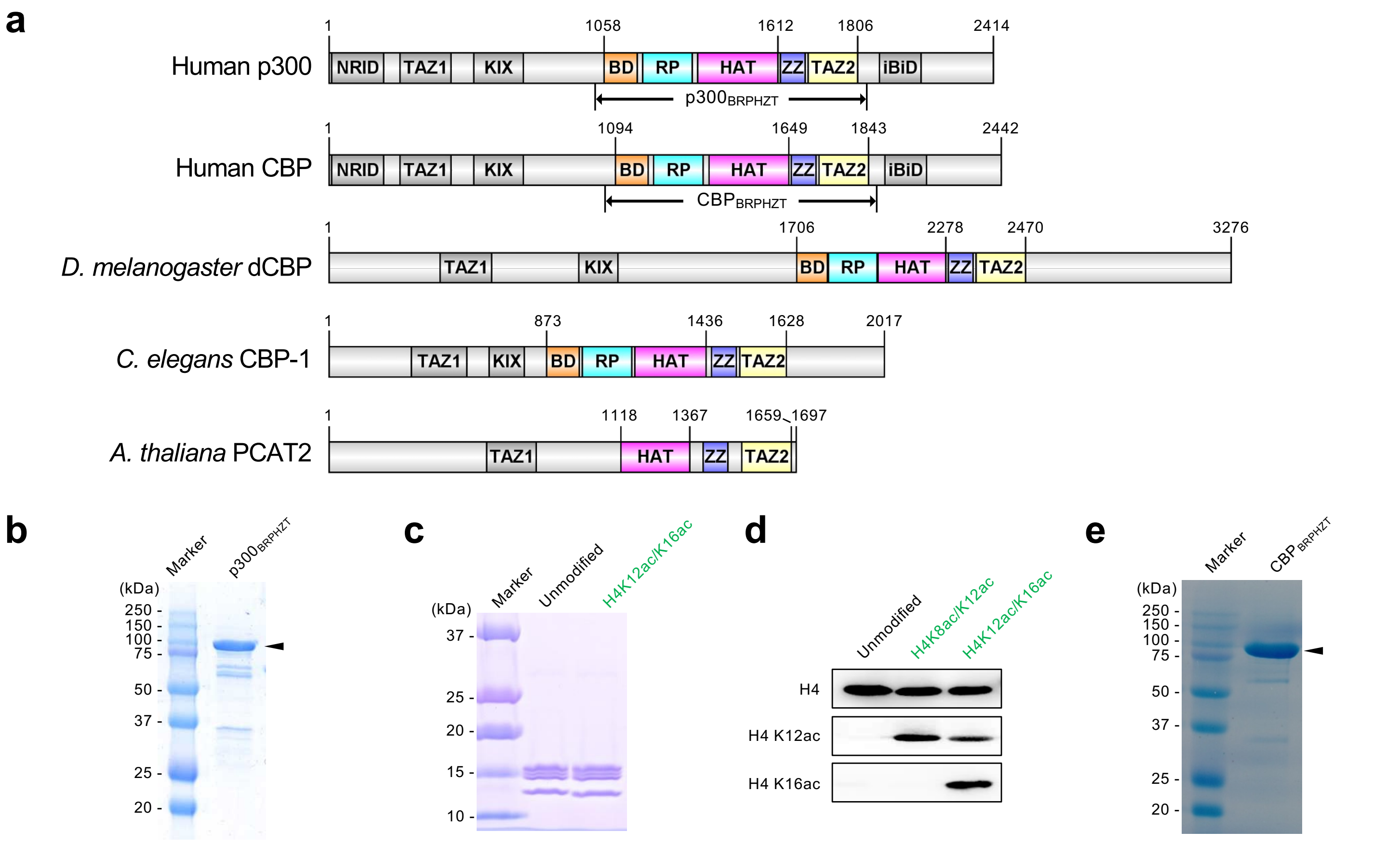

**Supplementary Figure 1 Enzymes and nucleosome substrates used in this study. a** Schematic representation of the domain architecture of metazoan p300/CREB-binding protein (CBP) homologs. NRID, nuclear receptor interaction domain; TAZ1, transcriptional adaptor zinc-finger domain 1; KIX, kinase-inducible domain of CREB-interacting domain; BD, bromodomain; RP, the RING and PHD zinc-fingers; HAT, histone acetyltransferase domain; ZZ, ZZ-type zinc-finger; TAZ2, transcriptional adaptor zinc-finger domain 2; and IBiD, IRF3-binding domain. The positions of the N- and C-termini and the start/end residues of the major domains are shown at the top of each scheme. The positions of the start/end residues of the human p300 and CBP constructs used in this study (*i.e.*, $p300_{BRPHZT}$ and $CBP_{BRPHZT}$) are shown at the bottom of each scheme. **b** A Coomassie Brilliant Blue (CBB)-stained *sodium* dodecyl sulfate polyacrylamide gel electrophoresis (SDS-PAGE) image of the $p300_{BRPHZT}$ protein. The size of each band of molecular weight markers is indicated on the left. The position of $p300_{BRPHZT}$ is indicated by an arrowhead on the right. **c** Preparation of the nucleosome containing K12/K16-di-acetylated H4. A CBB-stained SDS-PAGE image of the reconstituted nucleosomes is shown. The positions of acetyllysine introduced into histone H4 in the nucleosome are given in green at the top. **d** Immunoblotting of residue-specific histone H4 acetylation of the reconstituted nucleosomes. The positions of acetyllysine introduced into histone H4 in the nucleosome are given in green at the top. The residue-specific histone acetylation recognition antibody used is shown on the left. **e** A CBB-stained SDS-PAGE image of the $CBP_{BRPHZT}$ protein. The position of $CBP_{BRPHZT}$ is indicated by an arrowhead on the right.

### Supplementary Figure 2

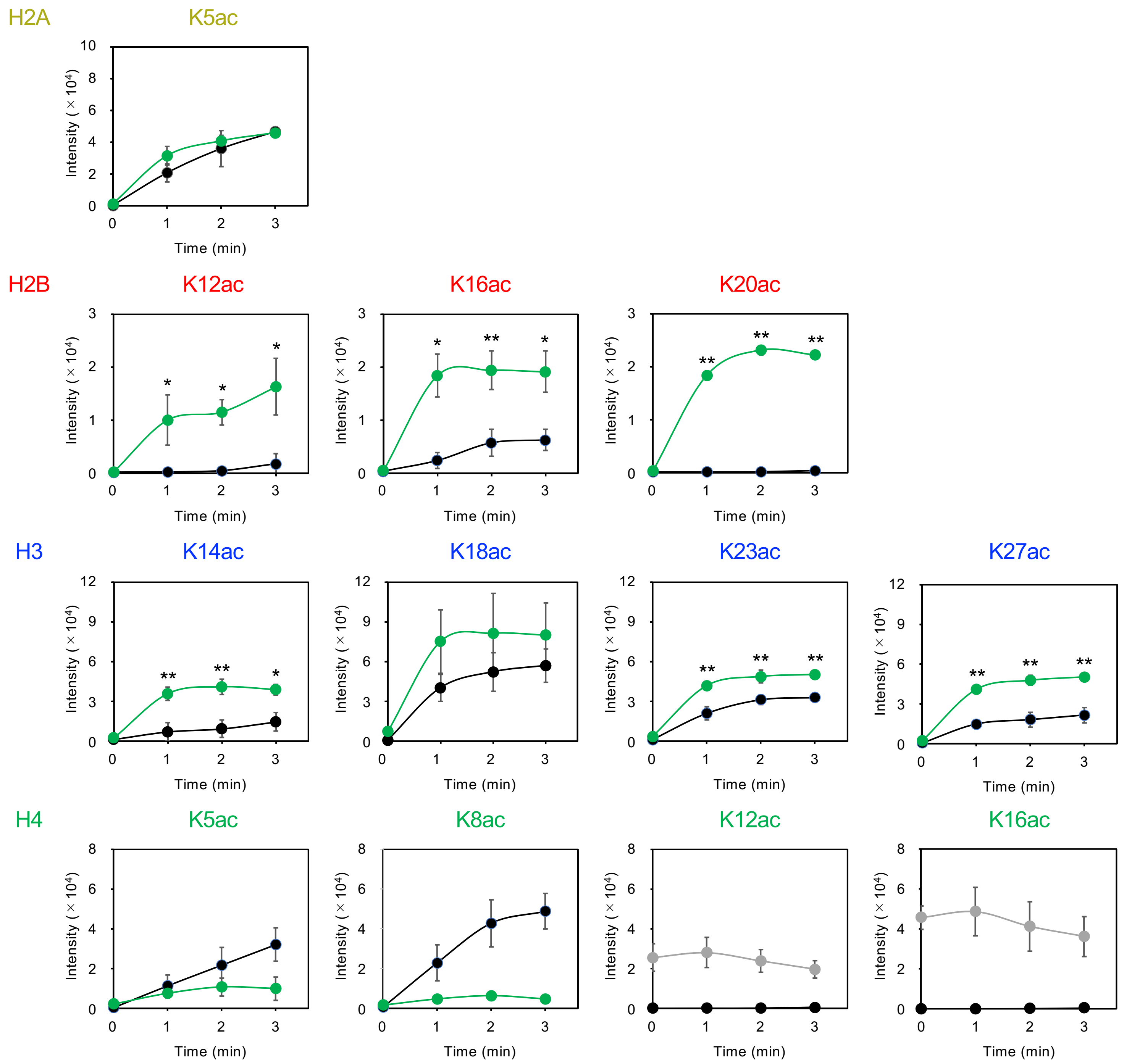

**Supplementary Figure 2 *In vitro* acetyltransferase activity of p300$_{\mathrm{BRPHZT}}$ toward an H4-di-acetylated nucleosome.** Residue-specific acetylation for each histone species detected by immunoblotting. The position of acetylation is shown above each panel. Black and green lines indicate the unmodified and the H4K12/K16-acetylated nucleosomes as substrates (1 µM), respectively. For pre-acetylated H4K12ac and H4K16ac residues, data with the H4K12/K16-acetylated nucleosome as substrate are shown as gray lines. The x-axis indicates the time course after the reaction in the presence of 1 µM p300$_{\mathrm{BRPHZT}}$ and 10 µM acetyl-CoA. The y-axis indicates the immunoblotting signal intensity. Means ± SD ($N = 3$). Statistical significance was assessed by a two-sample one-sided Welch's $t$-test for each time point ($^{*}P < 0.05$; $^{**}P < 0.01$). The alternative hypothesis is that the acetylated nucleosome is more acetylated by p300$_{\mathrm{BRPHZT}}$ than the unmodified nucleosome.

### Supplementary Figure 3

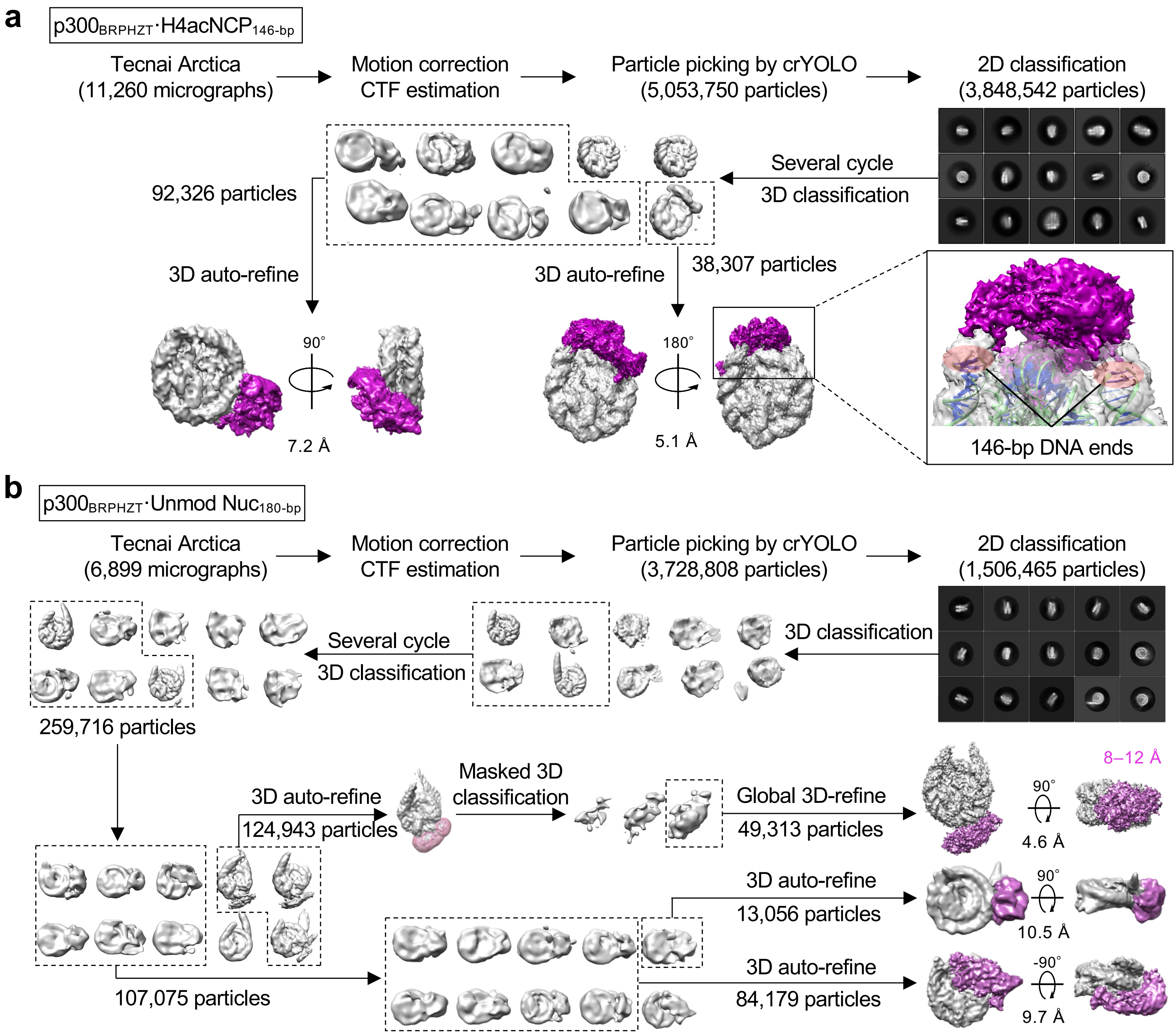

**Supplementary Figure 3 Cryo-electron microscopy workflow for proteins in complex with modified and unmodified nucleosomes. a** The processing pipeline for $p300_{BRPHZT}$ in complex with H4K12/K16-acetylated nucleosome using 146-bp double-stranded DNA ($H4acNCP_{146\text{-}bp}$). **b** The cryo-EM processing pipeline for $p300_{BRPHZT}$ with unmodified nucleosome using 180-bp double-stranded DNA (Unmod $Nuc_{180\text{-}bp}$). For both pipelines, the resolution was estimated by the gold standard Fourier shell correlation with the 0.143 criterion. Automated particle picking was performed with crYOLO. CTF, contrast transfer function.

### Supplementary Figure 4

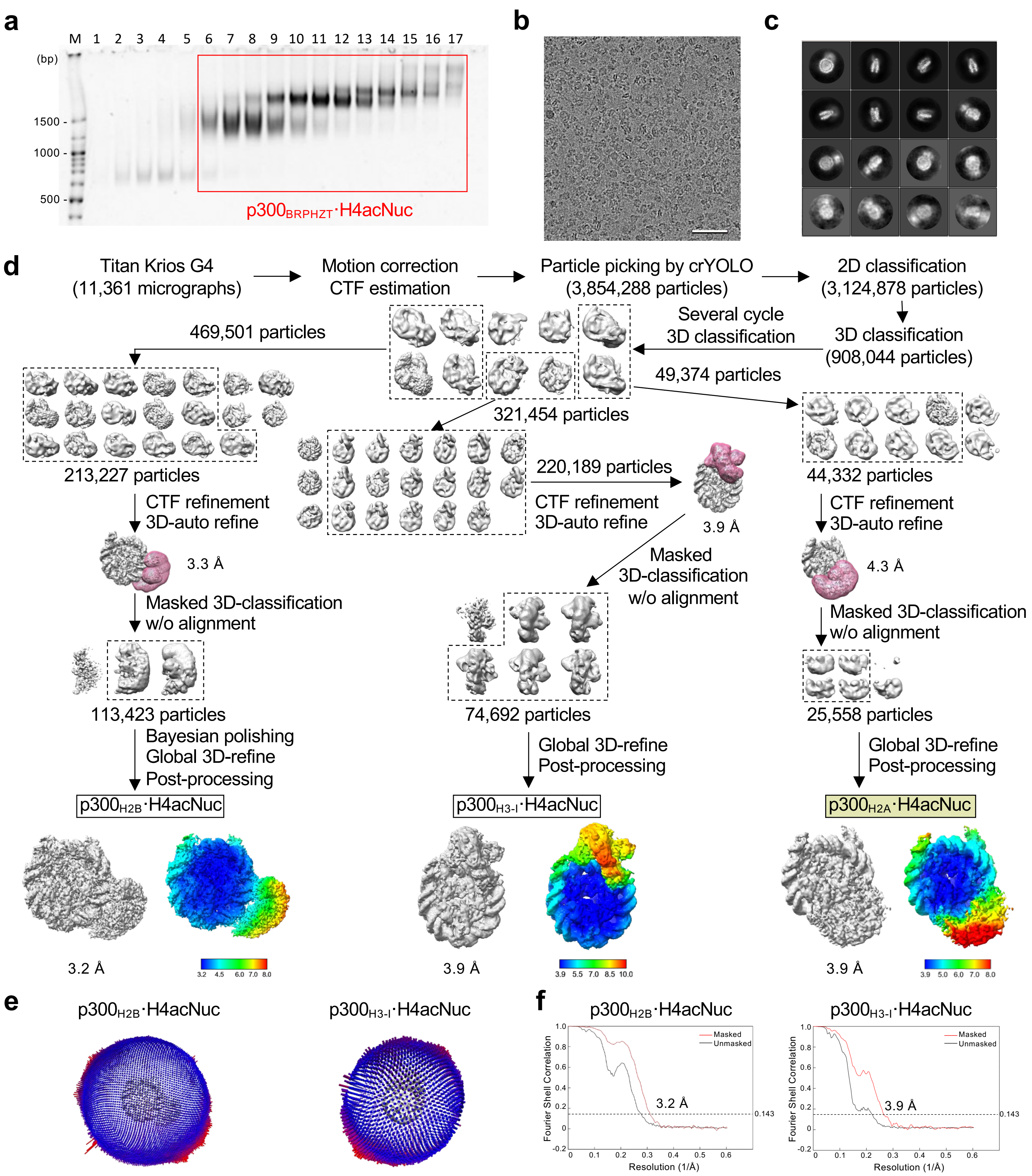

**Supplementary Figure 4 Sample preparation and cryo-electron microscopy structure determination of p300 protein in complex with reconstituted nucleosome (Titan Krios 300kV microscope). a** p300$_{BRPHZT}$ and reconstituted nucleosome (H4acNuc) were incubated and isolated by glycerol-gradient (10–30%) with crosslinking. Fractions 1–17 were analyzed by electrophoretic mobility shift assay. Bands are labeled on the left. Fractions 6–17 in the red box were used for cryo-electron microscopy. **b** Example cryo-electron micrograph. Scale bar, 50 nm. **c** Representative 2D class averages. **d** The processing pipeline for the p300$_{BRPHZT}$·H4acNuc complex. Local resolution (Å) was displayed on the sharpened full map. **e** Angular distribution of particle projections of the final reconstruction. **f** Gold-standard Fourier shell correlation curves of the final reconstitution. The resolution was estimated with the 0.143 criterion. Automated particle picking was performed with crYOLO. CTF, contrast transfer function.

### Supplementary Figure 5

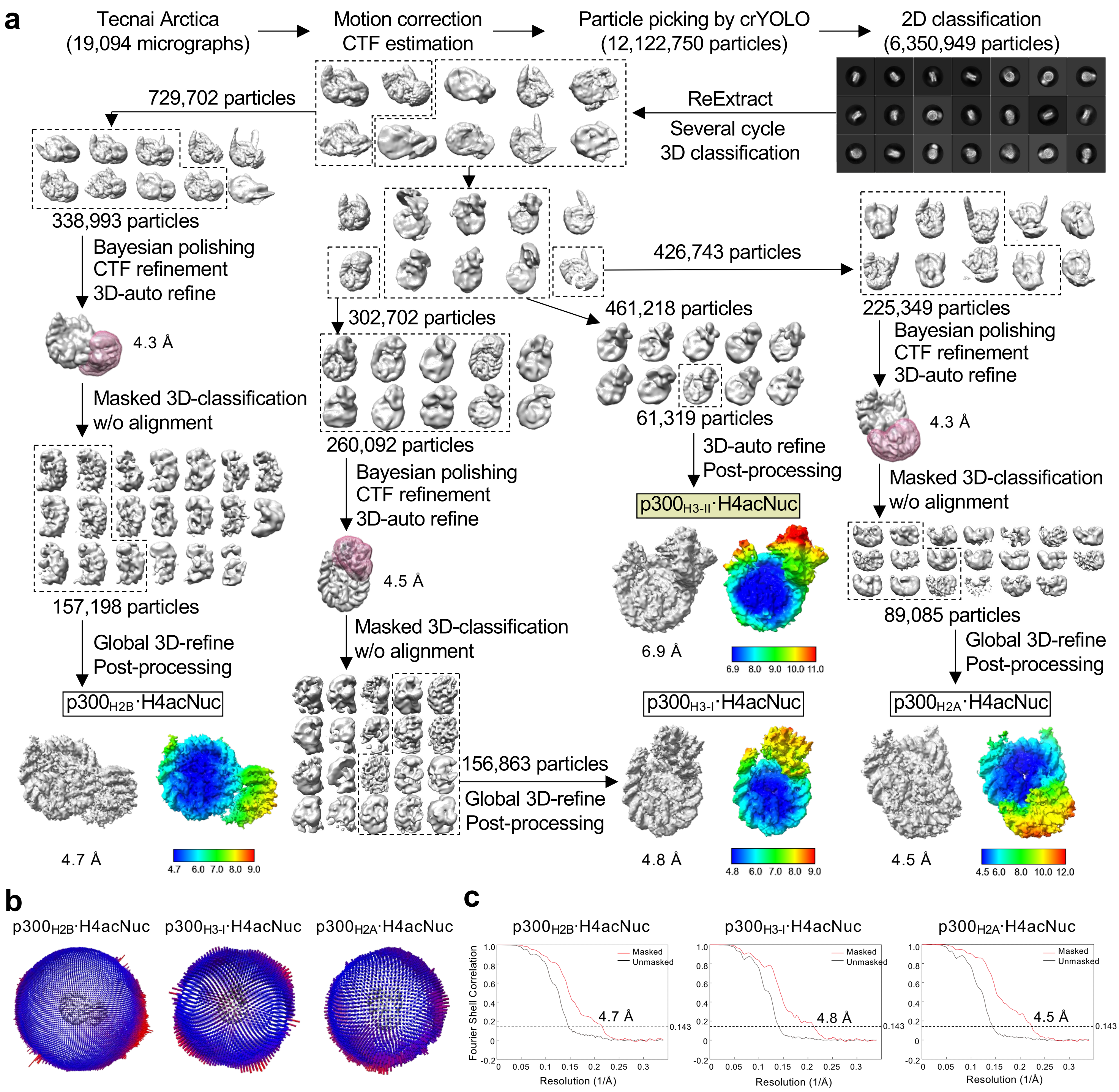

**Supplementary Figure 5 Sample preparation and cryo-electron microscopy structure determination of p300 protein in complex with reconstituted nucleosome (Tecnai Arctica 200kV microscope). a** Processing pipeline for the p300$_{BRPHZT}$·reconstituted nucleosome (H4acNuc) complex. Local resolution (Å) was displayed on the sharpened full map. **b** Angular distribution of particle projections of the final reconstruction. **c** Gold-standard Fourier shell correlation curves of final reconstitution. The resolution was estimated with the 0.143 criterion. Automated particle picking was performed with crYOLO. CTF, contrast transfer function.

### Supplementary Figure 6

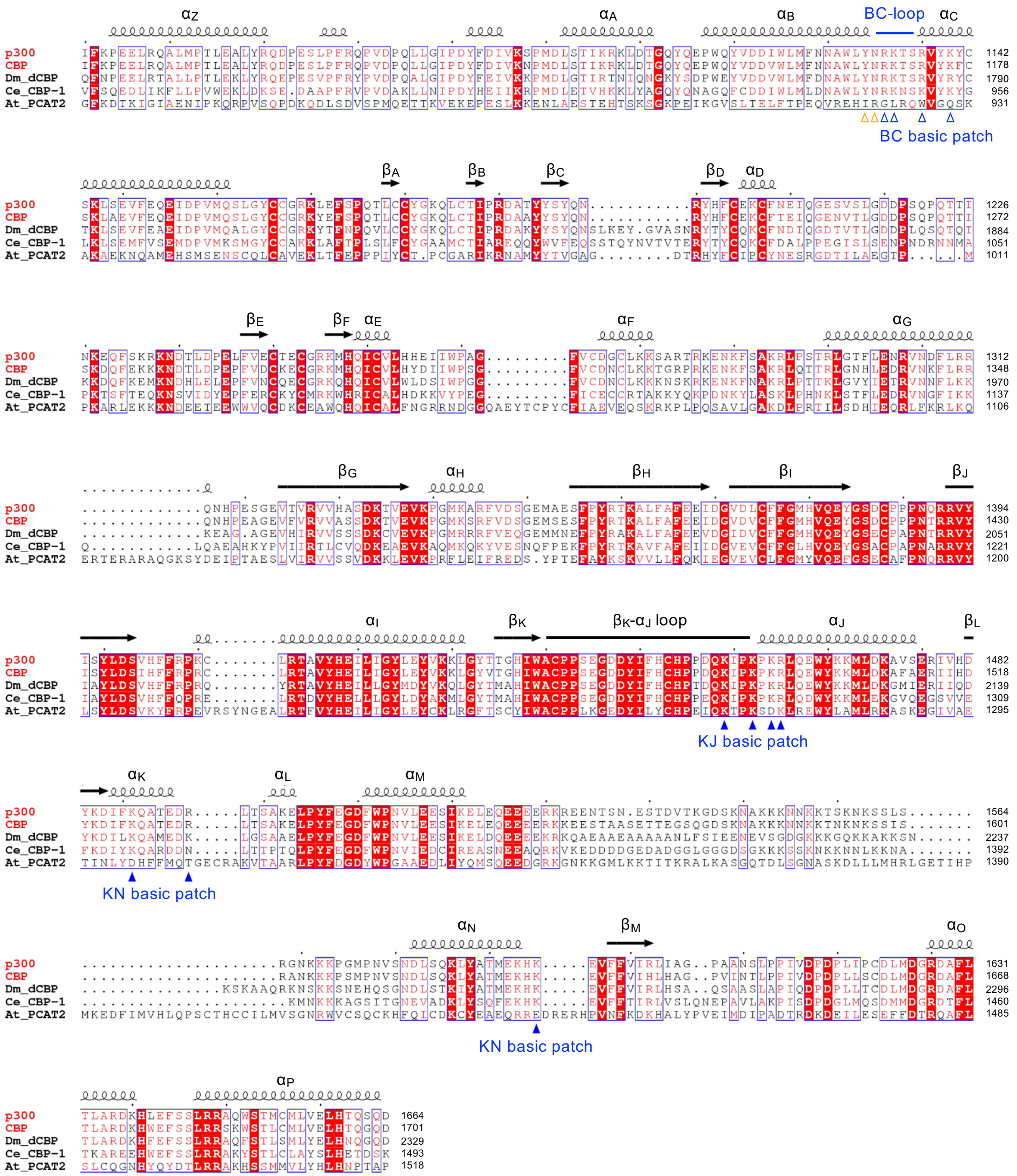

**Supplementary Figure 6 Sequence alignment of BRPH in p300/CBP homologs.** Homology between human p300 and CBP, *D. melanogaster* dCBP, *C. elegans* CBP-1, and *A. thaliana* PCAT2 is shown. Residue numbers on the C-terminal side of each row are shown on the right. Conserved or similar residues are shown in red and surrounded by blue boxes. The completely conserved residues are shown in white letters on a red background. The positions and numbers of α-helices and β-strands are indicated at the top of the alignment. The α-helices are numbered to match the bromodomain helix numbers ($\alpha Z$ to $\alpha C$). K/R residues comprising the two DNA-interactive basic patches in human p300 are indicated by filled blue arrowheads at the bottom (KJ basic patch: K1456, K1459, K1461, and R1462; KN basic patch: K1488, R1494, and K1592). Residues involved in the recognition of acetyllysine inside the bromodomain pocket (Y1131 and N1132) are indicated by open orange arrowheads. K/R residues comprising the third basic patch around the BC-loop of the bromodomain (BC basic patch; R1133, K1134, R1137, and K1140) are indicated by open blue arrowheads.

### Supplementary Figure 7

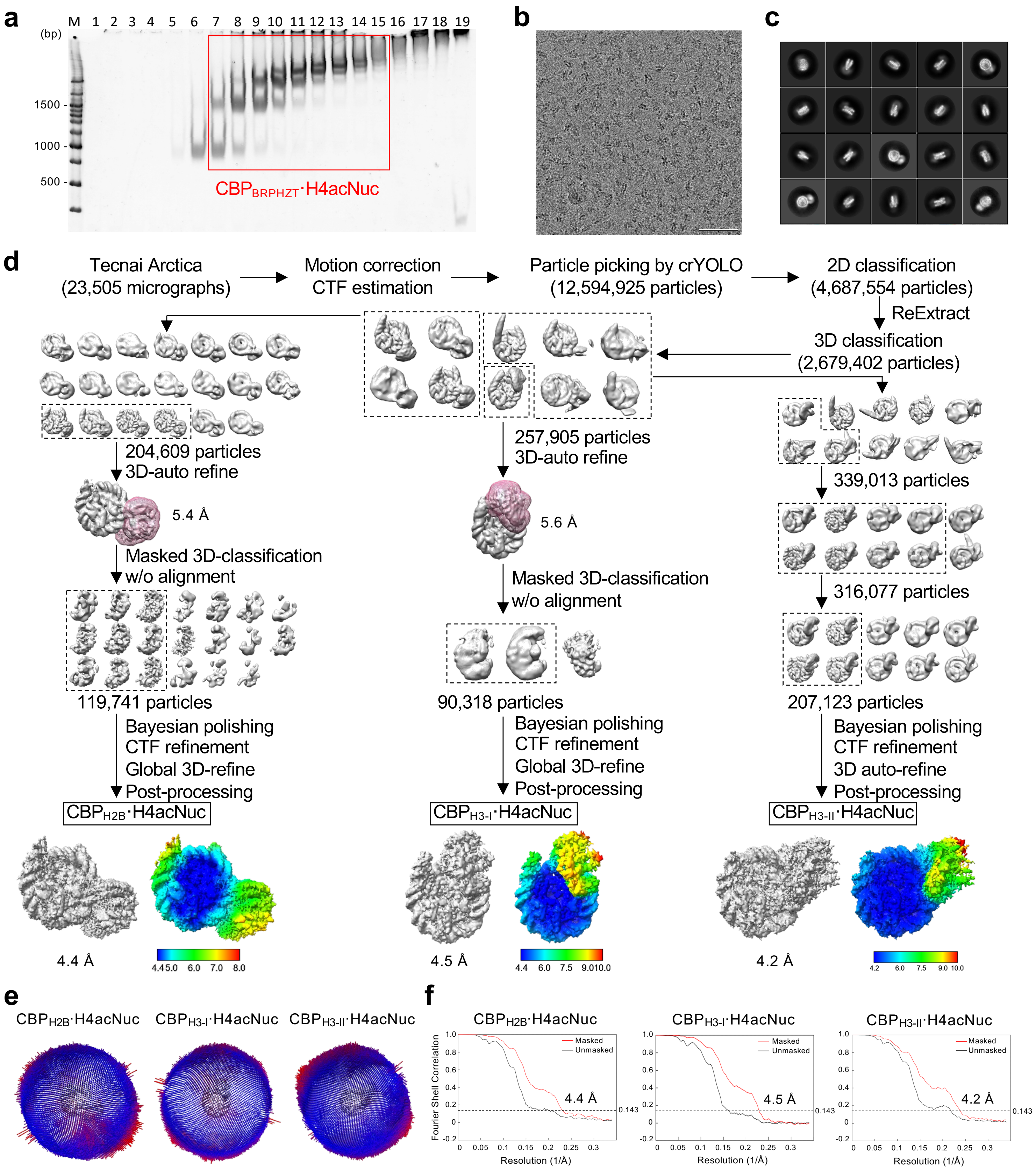

**Supplementary Figure 7 Sample preparation and cryo-electron microscopy structure determination of CREB-binding protein (CBP) in complex with reconstituted nucleosome. a** CBP$_{BRPHZT}$ and the reconstituted nucleosome (H4acNuc) were incubated and isolated by glycerol-gradient (10–40%) with crosslinking. Fractions 1–17 were analyzed by electrophoretic mobility shift assay. Bands are labeled on the left. Fractions 7–15 (red box) were used for cryo-electron microscopy. **b** Example cryo-electron micrograph. Scale bar, 50 nm. **c** Representative 2D class averages. **d** The microscopy processing pipeline for CBP$_{BRPHZT}$·H4acNuc complex. Local resolution (Å) is displayed on the sharpened full map. **e** Angular distribution of particle projections of the final reconstruction. **f** Gold-standard Fourier shell correlation curves of the final reconstitution. The resolution was estimated with the 0.143 criterion. Automated particle picking was performed with crYOLO. CTF, contrast transfer function.

### Supplementary Figure 8

| | p300$_{BRPH}$ (Titan Krios G4) | p300$_{BRPH}$ (Tecnai Arctica) | CBP$_{BRPH}$ (Tecnai Arctica) |
| --- | --- | --- | --- |
| H2B | #1 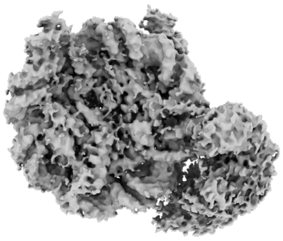 3.2 Å | #4 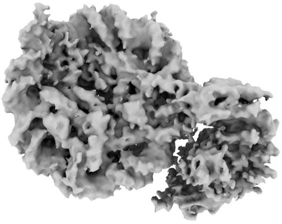 4.7 Å | #8 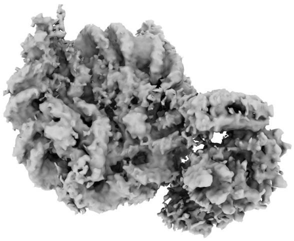 4.4 Å |
| H3 model I | #2 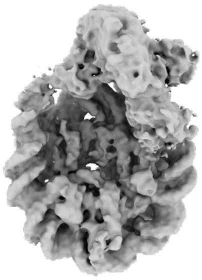 3.9 Å | #5 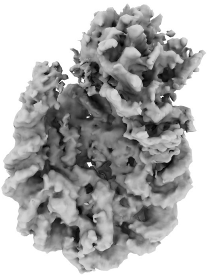 4.8 Å | #9 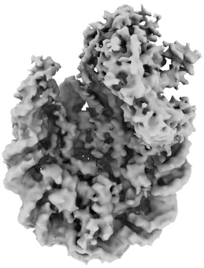 4.5 Å |
| H3 model II | NA | #6 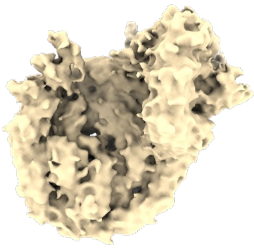 6.9 Å | #10 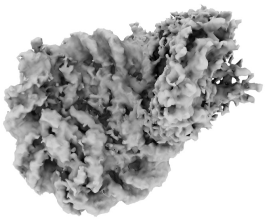 4.2 Å |
| H2A | #3 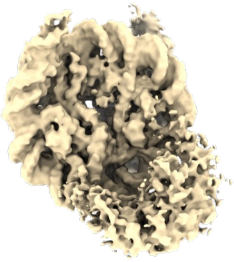 3.9 Å | #7 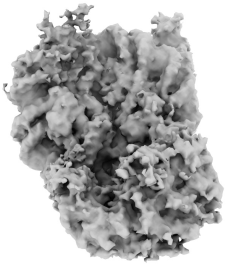 4.5 Å | NA |

**Supplementary Figure 8 All cryo-EM maps of the p300$_{BRPH}$/CBP$_{BRPH}$ complexed with H4acNuc.** The resolution was estimated by the gold standard Fourier shell correlation with the 0.143 criterion. Cryo-EM maps for which the atomic coordinates were determined are shown in dark gray. Those for which the atomic coordinates could not be determined are shown in light yellow. NA, not available.

### Supplementary Figure 9

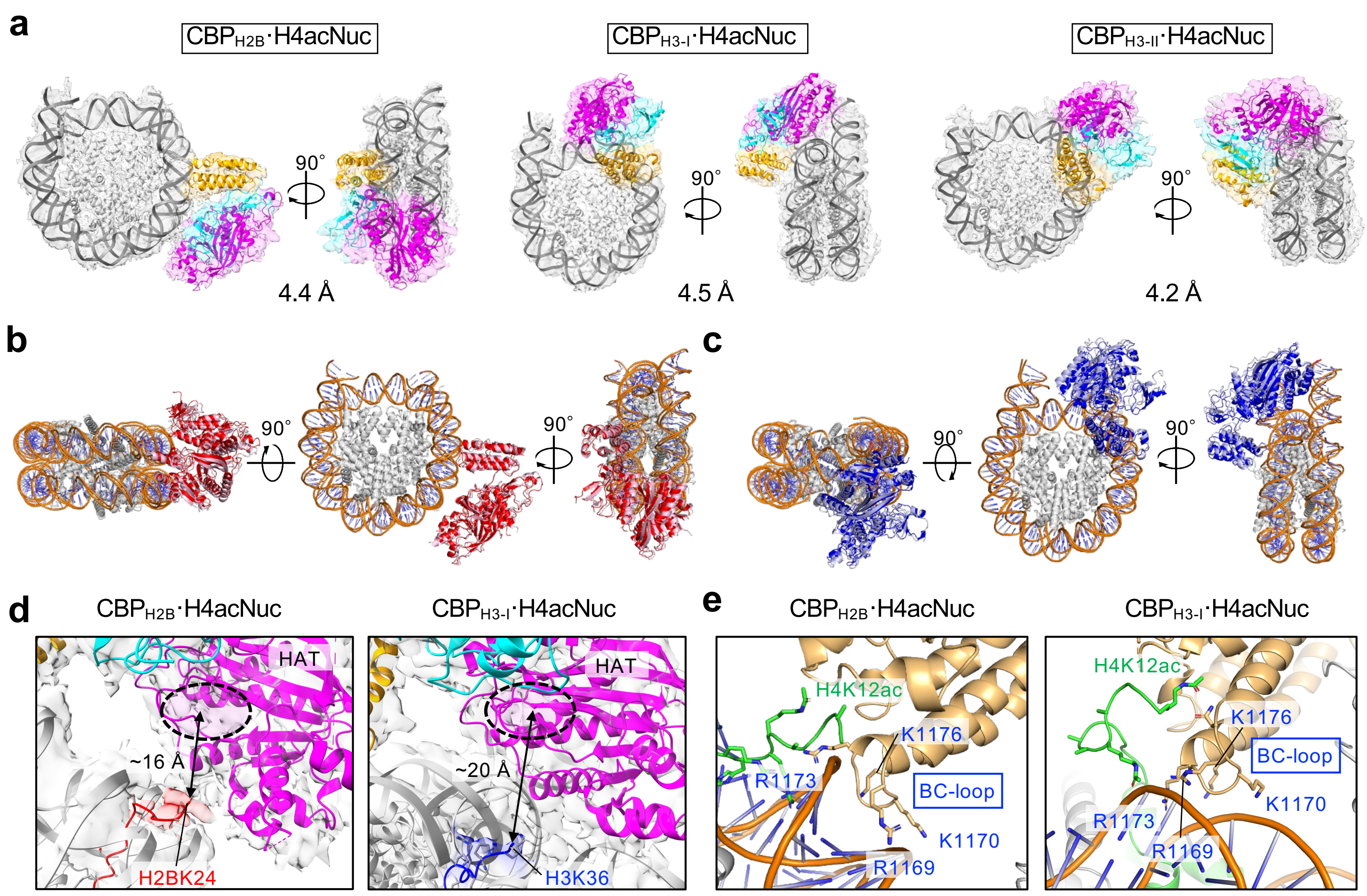

**Supplementary Figure 9 Cryo-electron microscopy structure in complex with reconstituted nucleosome. a** Various conformations of CBP$_{BRPH}$ and reconstituted nucleosome (H4acNuc) in cryo-electron microscopy (cryo-EM) maps and structural model. Left, CBP$_{H2B}$·H4acNuc (#8 in Supplementary Fig. 8); center, CBP$_{H3-I}$·H4acNuc (#9); right, CBP$_{H3-II}$·H4acNuc (#10). **b** Superimposition of CBP$_{H2B}$·H4acNuc (red) on p300$_{H2B}$·H4acNuc (pink). **c** Superimposition of CBP$_{H3-I}$·H4acNuc (blue) on p300$_{H3-I}$·H4acNuc (light blue). **d** Close-up views (#8 and #9) of the structural model and cryo-EM maps of H4K12acK16ac binding by BD of each complex structure. Color code: orange, CBP BD; cyan, CBP RP; magenta, CBP HAT; green, K12/K16-acetylated H4. **e** Close-up views (#8 and #9) of the interaction region by lysine and arginine in the BC-loop of the CBP of each complex structure.

### Supplementary Figure 10

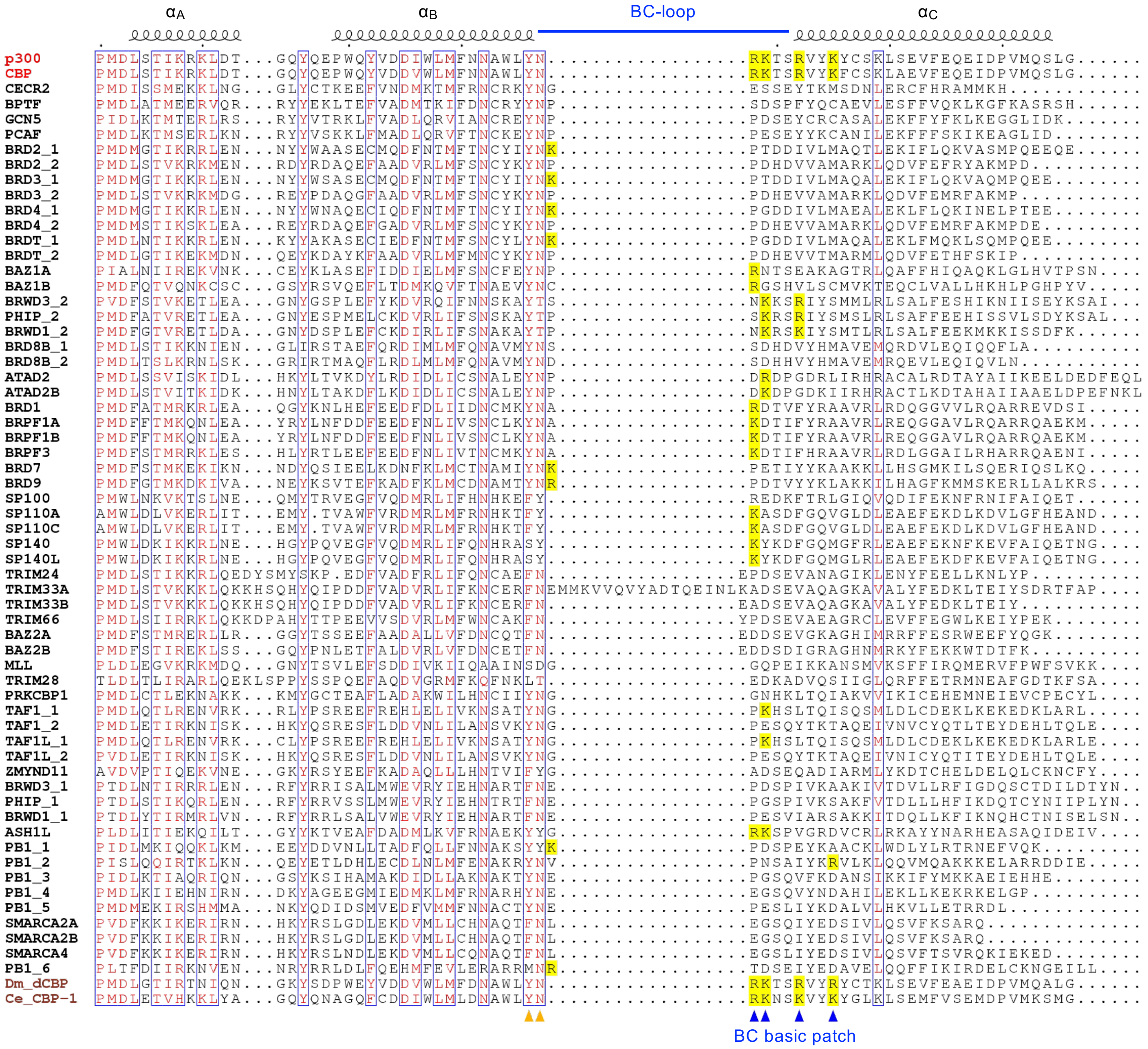

**Supplementary Figure 10 Sequence alignment around the BC-loop of the bromodomains.** The amino acid sequences of all 61 human bromodomains and the *D. melanogaster* and *C. elegans* p300/CBP homologs (*i.e.*, *Dm*_dCBP and *Ce*_CBP-1 shown in brown) are aligned. The positions of the BC-loop and three α-helices composing the bromodomain are shown on the top in black and blue, respectively. Protein names of human bromodomains other than p300 and CBP are shown in black on the left. Conserved or similar residues are shown in red and surrounded by blue boxes. Residues involved in the recognition of acetyllysine inside the bromodomain pocket (Y1131 and N1132) are indicated by filled orange arrowheads at the bottom. K/R residues comprising the basic patch around the BC-loop of the bromodomain (BC basic patch; R1133, K1134, R1137, and K1140 of human p300) are indicated by a yellow background and filled blue arrowheads at the bottom.

### Supplementary Figure 11

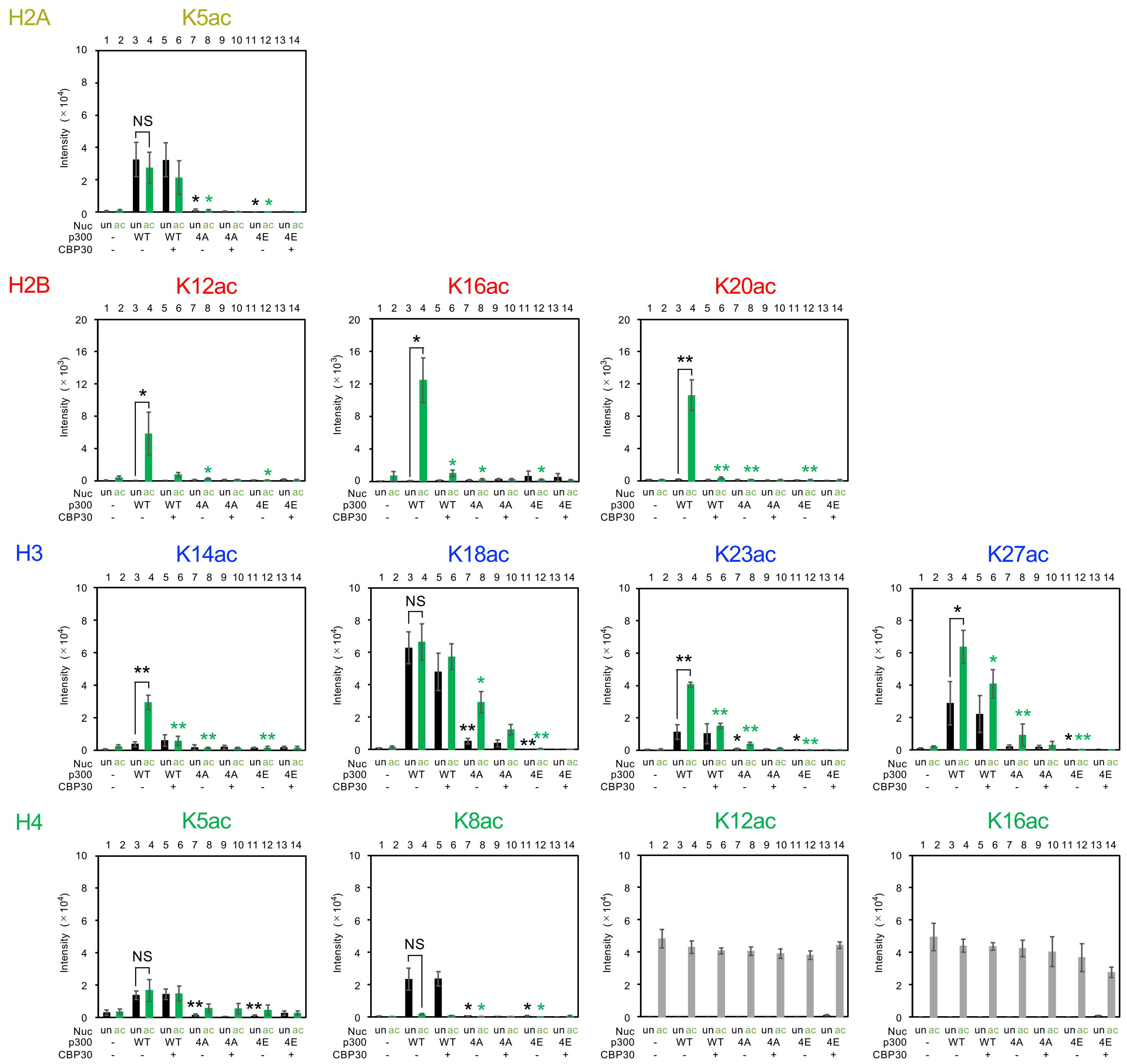

**Supplementary Figure 11 Effect of p300 mutations on its acetyltransferase activity toward the H4-di-acetylated nucleosome.** Residue-specific histone acetylation detected by immunoblotting for each histone species. The position of acetylation is shown above each panel. Nucleosome (Nuc): un, unmodified; ac (green), H4K12/K16-acetylated. p300: WT, wild-type $\text{p300}_{\text{BRPHZT}}$; 4A, $\text{p300}_{\text{BRPHZT}}$ with mutations of R1133A, K1134A, R1137A, and K1140A; 4E, $\text{p300}_{\text{BRPHZT}}$ with mutations of R1133E, K1134E, R1137E, and K1140E; and CBP30: -, none; +, 10 μM. The y-axis indicates the immunoblotting signal intensity at 1 min after the reaction. Means ± SD ($N = 3$). For pre-acetylated H4K12ac and H4K16ac residues, data with the H4K12/K16-acetylated nucleosome as substrate are shown as gray bars. Statistical significance was assessed by a two-sample one-sided Welch's $t$-test (NS, $P \geq 0.05$; $^{*}P < 0.05$; $^{**}P < 0.01$). The alternative hypothesis is as follows: lane 4, increase vs. lane 3; lanes 5, 7, and 11, decrease vs. lane 3; lanes 6, 8, and 12, decrease vs. lane 4).

### Supplementary Figure 12

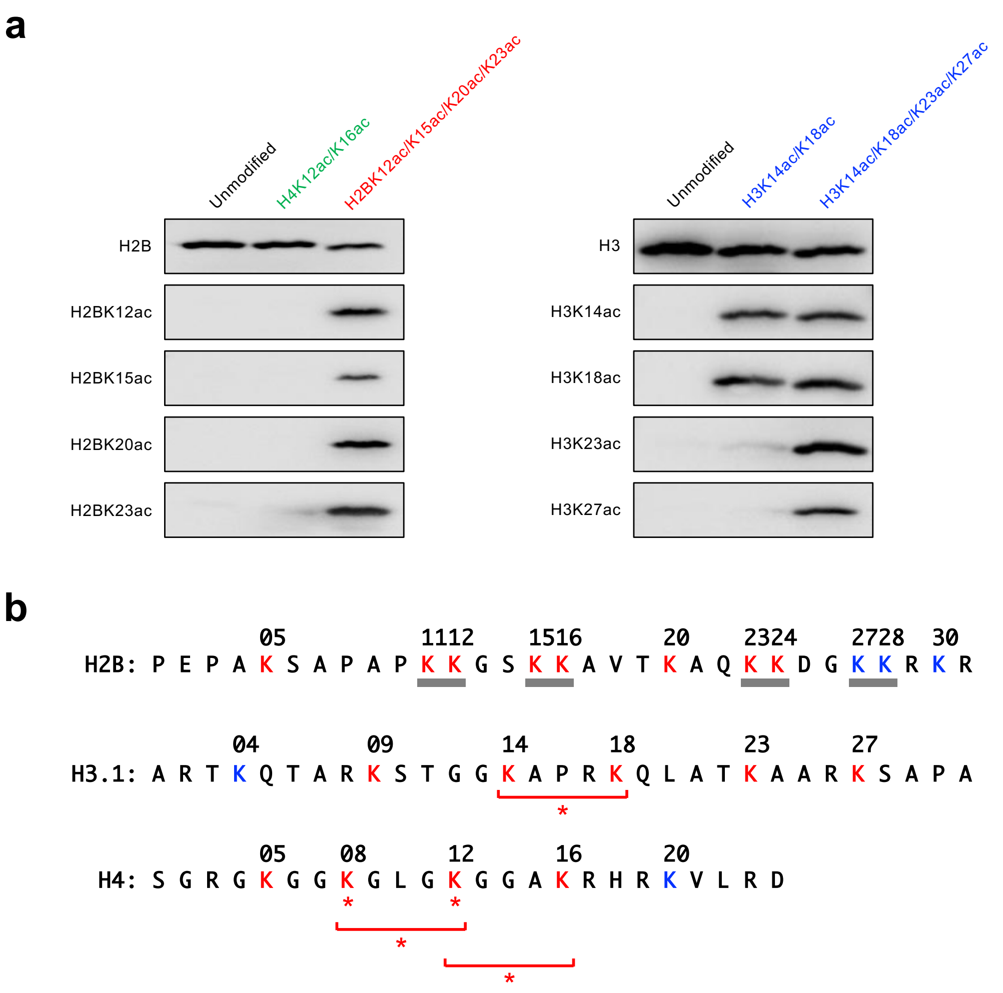

**Supplementary Figure 12 Lysine acetylation in the histone N-terminal tails (NTs). a** Immunoblotting of residue-specific histone acetylation of the reconstituted nucleosomes. Left, H4-di-acetylated or H2B-tetra-acetylated nucleosome; right, H3-di-acetylated or tetra-acetylated nucleosomes. The position(s) of acetyllysine introduced into histone H4, H2B, or H3 in the nucleosome are shown above each image. The residue-specific histone acetylation recognition antibody used is shown to the left of each image. **b** Amino acid sequence of human histone NTs. Histone names are shown on the left. The type of H2B is 1-J. Lysine (K) residues that p300 acetylates are indicated in red, and those it does not acetylate[15] in blue. Sequences with contiguous lysine residues (KK) are indicated by gray bars at the bottom. Kac residues or combinations of Kac residues to which p300 bromodomain binds are indicated by asterisks and underlines, respectively. A binding threshold was applied at a $K_D$ of ~100 µM[13].

### Supplementary Figure 13

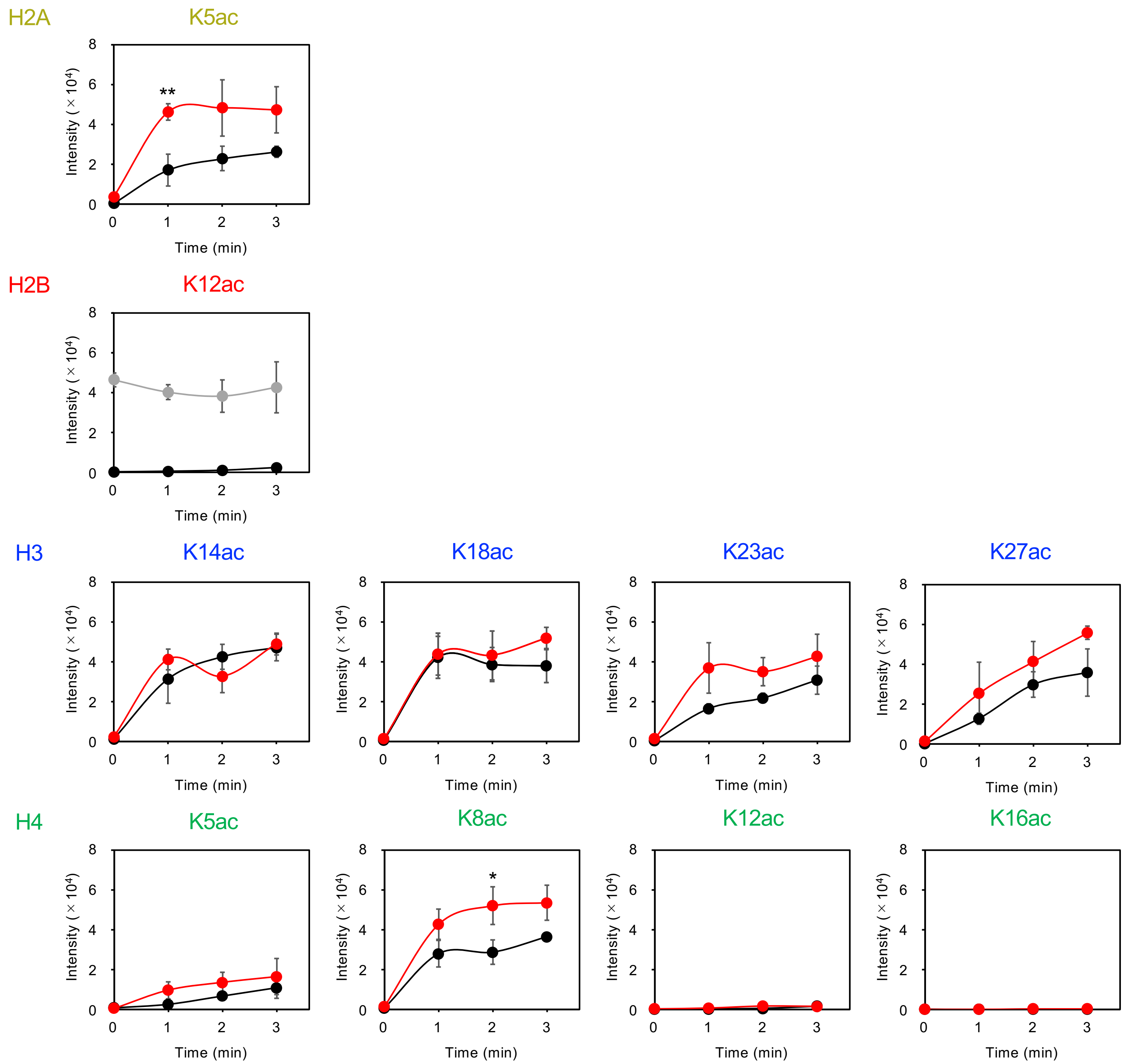

**Supplementary Figure 13 *In vitro* acetyltransferase activity of p300$_{BRPHZT}$ toward an H2B-tetra-acetylated nucleosome.** Residue-specific histone acetylation detected by immunoblotting for each histone species. The position of acetylation is shown above each panel. Black and red lines indicate the unmodified- and the H2BK12/K15/K20/K23-acetylated nucleosomes as substrates (1 μM), respectively. For pre-acetylated H2BK12ac residue, data with the H2BK12/K15/K20/K23-acetylated nucleosome as substrate is shown as gray line. The x-axis indicates the time course after the reaction in the presence of 1 μM p300$_{BRPHZT}$ and 10 μM acetyl-CoA. The y-axis indicates the immunoblotting signal intensity. Means ± SD ($N = 3$). Statistical significance was assessed by a two-sample one-sided Welch's $t$-test for each time point ($^{*}P < 0.05$; $^{**}P < 0.01$). The alternative hypothesis is that the acetylated nucleosome is more acetylated by p300$_{BRPHZT}$ than the unmodified nucleosome.

### Supplementary Figure 14

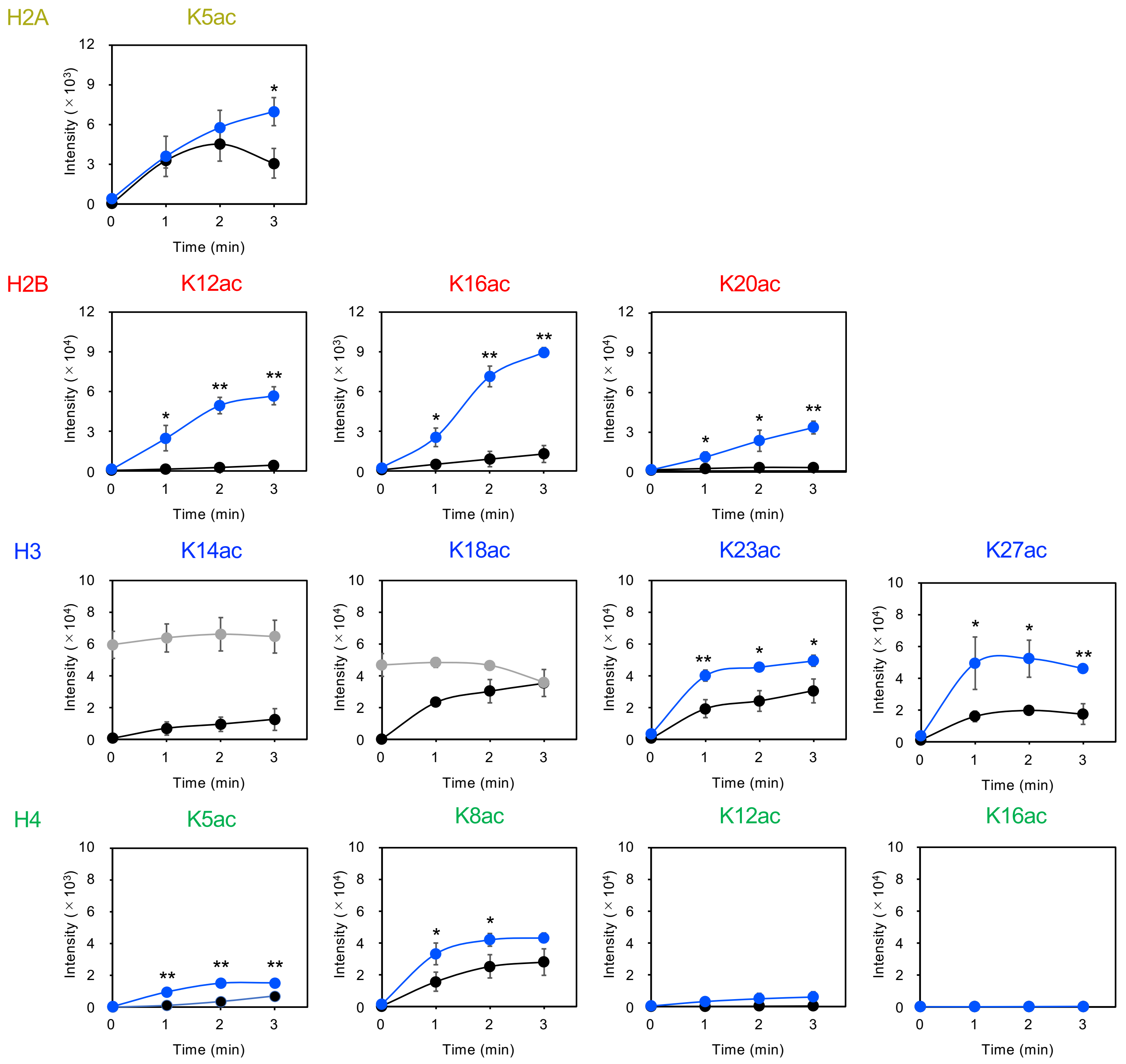

**Supplementary Figure 14 *In vitro* acetyltransferase activity of p300$_{BRPHZT}$ toward an H3-di-acetylated nucleosome.** Residue-specific histone acetylation detected by immunoblotting for each histone species. The position of acetylation is shown above each panel. Black and blue lines indicate the unmodified- and the H3K14/K18-acetylated nucleosomes as substrates (1 μM), respectively. For pre-acetylated H3K14ac and H3K18ac residues, data with the H3K14/K18-acetylated nucleosome as substrate are shown as gray lines. The x-axis indicates the time course after the reaction in the presence of 1 μM p300$_{BRPHZT}$ and 10 μM acetyl-CoA. The y-axis indicates the immunoblotting signal intensity. Means ± SD ($N = 3$). Statistical significance was assessed by a two-sample one-sided Welch's *t*-test for each time point ($^{*}P < 0.05$; $^{**}P < 0.01$). The alternative hypothesis is that the acetylated nucleosome is more acetylated by p300$_{BRPHZT}$ than the unmodified nucleosome.

### Supplementary Figure 15

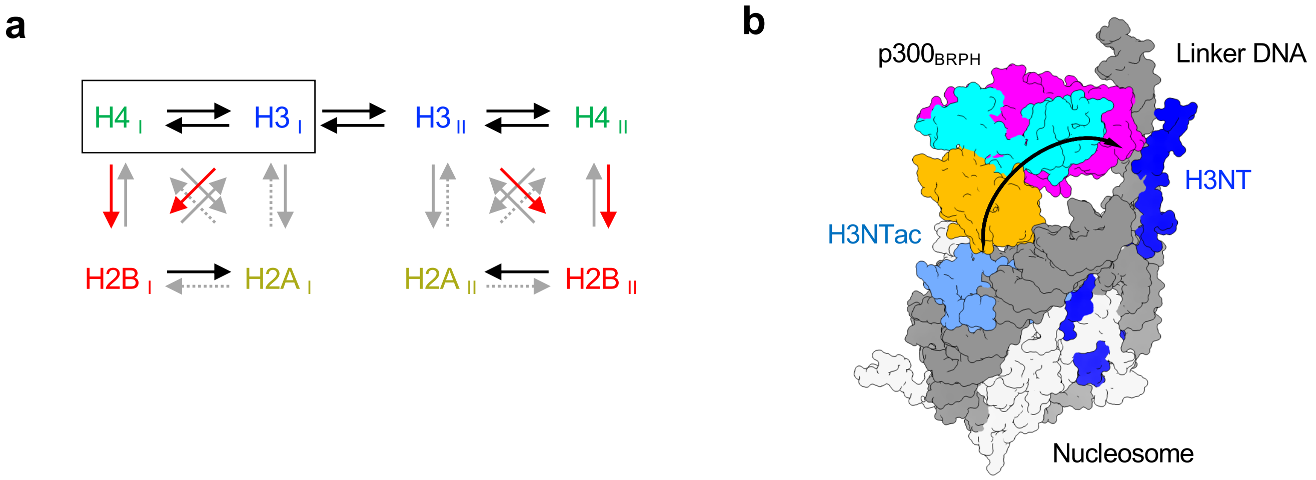

**Supplementary Figure 15 Flow of acetylation information propagated by p300. a** Summary of 'read/write' flow of histone N-terminal tail acetylation by p300. Each of the four pairs of histones in the nucleosome is shown schematically as I and II. Arrows indicate the 'read' → 'write' direction of histone acetylation and are classified as follows: red, high signal-to-noise ratio signalling; black, moderate signalling; gray, low or no signalling; dotted gray, not determined. The H3-H4 dimer, which could be derived from the parental histone octamer, is schematically enclosed. **b** Hypothetical model in which $p300_{BRPH}$ 'reads/writes' Kac between the H3NT pair. $p300_{BRPH}$ can bind to the nucleosome having linker DNA at both ends, with its bromodomain 'reading' one of a pair of H3 N-terminal tails and its catalytic center, the histone acetyltransferase domain, simultaneously 'writing' Kac to the other H3 N-terminal tail. Color code of $p300_{BRPH}$: orange, bromodomain; cyan, the RING and PHD zinc-fingers; magenta, histone acetyltransferase domain. Nucleosomes, linker DNA, and histone H3 are colored light gray, dark gray, and blue (one pale blue and the other dark blue), respectively. The black arrow indicates 'read' → 'write' direction.

### Supplementary Figure 16

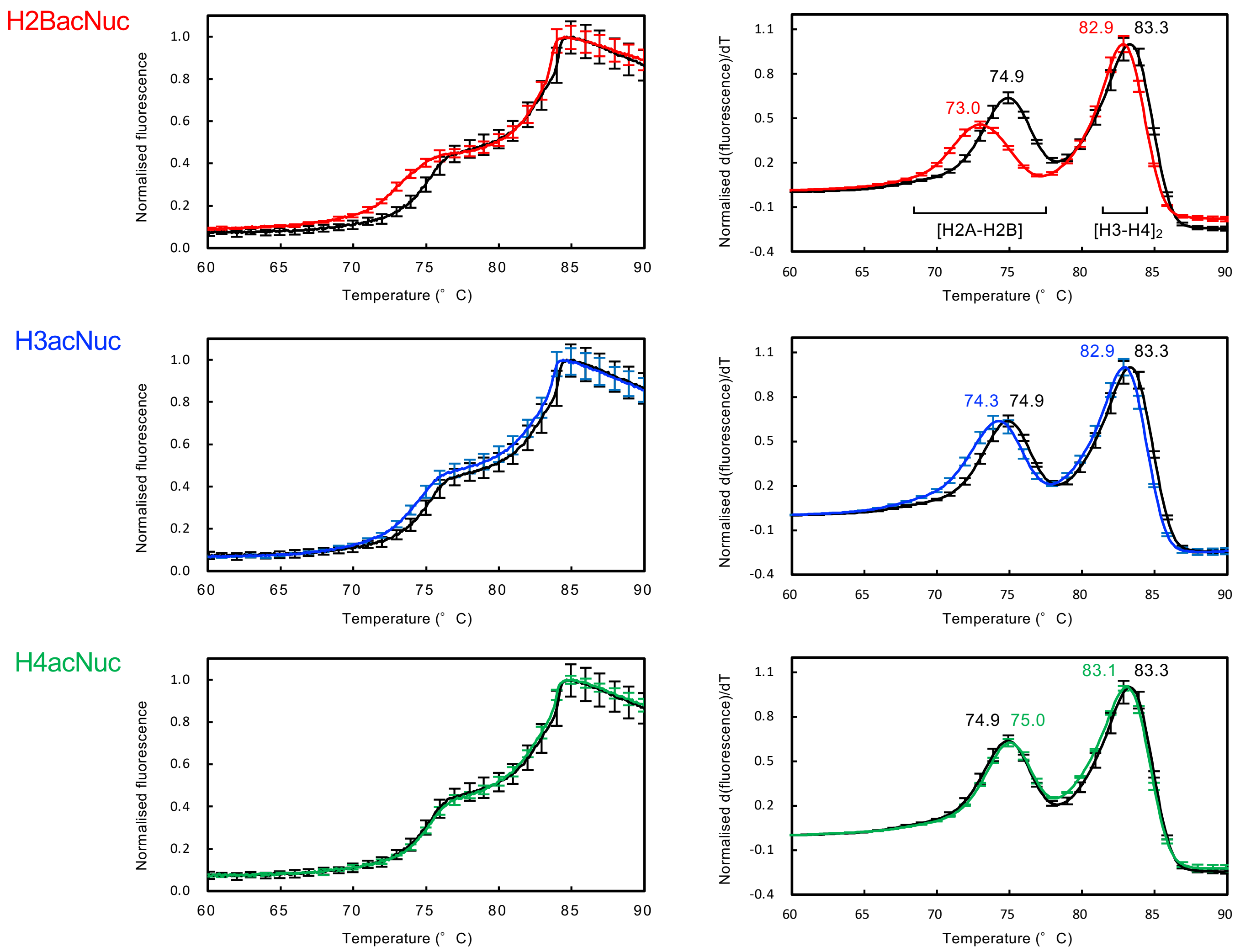

**Supplementary Figure 16 Thermal stability assay of acetylated nucleosomes.** Mean values of the thermal denaturation curves from 60.0 to 90.0 °C for fluorescence intensity (left) and derivative fluorescence intensity (right) are plotted. The black line indicates the unmodified nucleosome. The colored line indicates residue-specific acetylated nucleosome in the following color code: red, H2BK12/K15/K20/K23-acetylated; blue, H3K14/K18-acetylated; and green, H4K12/K16-acetylated. In the top right panel, the temperature at which the H2A-H2B dimer or the H3-H4 tetramer dissociates from the nucleosome is shown at the bottom. Means ± SD ($N = 3$).

#### Supplementary Table 1 Cryo-electron microscopy data collection, refinement, and validation statistics

| | #1 $p300_{H2B}$·H4acNuc (EMD-34588, PDB 8HAG) | #2 $p300_{H3-I}$·H4acNuc (EMD-34589, PDB 8HAH) | #3 $p300_{H2A}$·H4acNuc (EMD-34590) | #4 $p300_{H2B}$·H4acNuc (EMD-34591, PDB 8HAI) | #5 $p300_{H3-I}$·H4acNuc (EMD-34592, PDB 8HAJ) | #6 $p300_{H3-II}$·H4acNuc (EMD-34593) | #7 $p300_{H2A}$·H4acNuc (EMD-34594, PDB 8HAK) |
|---|---|---|---|---|---|---|---|
| **Data collection and processing** | | | | | | | |
| Microscope | Titan Krios G4 | Titan Krios G4 | Titan Krios G4 | Tecnai Arctica | Tecnai Arctica | Tecnai Arctica | Tecnai Arctica |
| Magnification | 105,000 | 105,000 | 105,000 | 23,500 | 23,500 | 23,500 | 23,500 |
| Voltage (kV) | 300 | 300 | 300 | 200 | 200 | 200 | 200 |
| Electron exposure (e–/Å$^2$) | 50 | 50 | 50 | 50 | 50 | 50 | 50 |
| Defocus range (μm) | -0.8– -2.0 | -0.8– -2.0 | -0.8– -2.0 | -0.9– -1.7 | -0.9– -1.7 | -0.9– -1.7 | -0.9– -1.7 |
| Pixel size (Å) | 0.829 | 0.829 | 0.829 | 1.47 | 1.47 | 1.47 | 1.47 |
| Symmetry imposed | C1 | C1 | C1 | C1 | C1 | C1 | C1 |
| Initial particle images (no.) | 3,854,288 | 3,854,288 | 3,854,288 | 12,122,750 | 12,122,750 | 12,122,750 | 12,122,750 |
| Final particle images (no.) | 113,423 | 74,692 | 25,558 | 157,198 | 156,863 | 61,319 | 89,085 |
| Map resolution (Å) | 3.2 | 3.9 | 3.9 | 4.7 | 4.8 | 6.9 | 4.5 |
| FSC threshold | 0.143 | 0.143 | 0.143 | 0.143 | 0.143 | 0.143 | 0.143 |
| Map resolution range (Å) | 3.1–8.4 | 3.5–10.6 | 3.6–10.3 | 4.3–11.3 | 3.1–9.3 | 3.1–9.3 | 4.1–12.7 |
| **Refinement** | | | | | | | |
| Initial model used (PDB code) | 1KX3, 6GYR | 1KX3, 6GYR | | 1KX3, 6GYR | 1KX3, 6GYR | | 1KX3, 6GYR |
| Model resolution (Å) | 3.6 | 8.6 | | 6.0 | 6.7 | | 6.0 |
| FSC threshold | 0.5 | 0.5 | | 0.5 | 0.5 | | 0.5 |
| Map sharpening *B* factor (Å$^2$) | -10 | -30 | | -180 | -119 | | -90 |
| Model-map scores | | | | | | | |
| CC (mask) | 0.78 | 0.62 | | 0.72 | 0.76 | | 0.75 |
| CC (box) | 0.85 | 0.83 | | 0.83 | 0.88 | | 0.87 |
| CC (peaks) | 0.70 | 0.42 | | 0.68 | 0.69 | | 0.69 |
| CC (volume) | 0.76 | 0.56 | | 0.73 | 0.74 | | 0.74 |
| Model composition | | | | | | | |
| Non-hydrogen atoms | 16,367 | 17,096 | | 16,547 | 17,214 | | 16,342 |
| Protein residues | 1284 | 1287 | | 1310 | 1305 | | 1285 |
| Nucleotides | 294 | 328 | | 294 | 326 | | 292 |
| R.m.s. deviations | | | | | | | |
| Bond lengths (Å) | 0.003 | 0.003 | | 0.003 | 0.003 | | 0.003 |
| Bond angles (°) | 0.547 | 0.539 | | 0.566 | 0.592 | | 0.641 |
| **Validation** | | | | | | | |
| MolProbity score | 1.54 | 1.59 | | 1.55 | 1.63 | | 1.67 |
| Clashscore | 10.55 | 8.69 | | 10.75 | 11.31 | | 12.75 |
| Poor rotamers (%) | 0.00 | 0.00 | | 0.00 | 0.00 | | 0.00 |
| Ramachandran plot | | | | | | | |
| Favored (%) | 98.33 | 97.39 | | 98.52 | 98.04 | | 97.77 |
| Allowed (%) | 1.67 | 2.61 | | 1.48 | 1.96 | | 2.33 |
| Disallowed (%) | 0.00 | 0.00 | | 0.00 | 0.00 | | 0.00 |

**Supplementary Table 1 (continued) Cryo-electron microscopy data collection, refinement, and validation statistics**

| | #8 $CBP_{H2B}$·H4acNuc (EMD-34595, PDB 8HAL) | #9 $CBP_{H3-I}$·H4acNuc (EMD-34596, PDB 8HAM) | #10 $CBP_{H3-II}$·H4acNuc (EMD-34597, PDB 8HAN) |
|---|---|---|---|
| **Data collection and processing** | | | |
| Microscope | Tecnai Arctica | Tecnai Arctica | Tecnai Arctica |
| Magnification | 23,500 | 23,500 | 23,500 |
| Voltage (kV) | 200 | 200 | 200 |
| Electron exposure (e–/Å$^2$) | 50 | 50 | 50 |
| Defocus range (μm) | -0.9– -1.7 | -0.9– -1.7 | -0.9– -1.7 |
| Pixel size (Å) | 1.47 | 1.47 | 1.47 |
| Symmetry imposed | C1 | C1 | C1 |
| Initial particle images (no.) | 12,594,925 | 12,594,925 | 12,594,925 |
| Final particle images (no.) | 119,741 | 90,318 | 207,123 |
| Map resolution (Å) | 4.4 | 4.5 | 4.2 |
| FSC threshold | 0.143 | 0.143 | 0.143 |
| Map resolution range (Å) | 4.0–8.7 | 4.1–12.1 | 3.9–12.4 |
| **Refinement** | | | |
| Initial model used (PDB code) | 1KX3, 5U7G | 1KX3, 5U7G | 1KX3, 5U7G |
| Model resolution (Å) | 6.1 | 4.5 | 6.6 |
| FSC threshold | 0.5 | 0.5 | 0.5 |
| Map sharpening *B* factor (Å$^2$) | -50 | -100 | -60 |
| Model-map scores | | | |
| CC (mask) | 0.73 | 0.77 | 0.72 |
| CC (box) | 0.85 | 0.84 | 0.83 |
| CC (peaks) | 0.67 | 0.72 | 0.67 |
| CC (volume) | 0.73 | 0.78 | 0.72 |
| Model composition | | | |
| Non-hydrogen atoms | 16,585 | 17,177 | 16,180 |
| Protein residues | 1292 | 1288 | 1261 |
| Nucleotides | 302 | 330 | 294 |
| R.m.s. deviations | | | |
| Bond lengths (Å) | 0.003 | 0.003 | 0.003 |
| Bond angles (°) | 0.619 | 0.555 | 0.520 |
| **Validation** | | | |
| MolProbity score | 1.60 | 1.50 | 1.45 |
| Clashscore | 12.30 | 9.55 | 8.34 |
| Poor rotamers (%) | 0.00 | 0.00 | 0.00 |
| Ramachandran plot | | | |
| Favored (%) | 98.26 | 98.57 | 98.86 |
| Allowed (%) | 1.74 | 1.43 | 1.14 |
| Disallowed (%) | 0.00 | 0.00 | 0.00 |

**Supplementary Table 2 Binding analysis between p300 bromodomain and nucleosomes, measured by microscale thermophoresis**

| $p300_{BRP}$ | Nucleosome | CBP30 | $K_{1/2}$ (nM) |
|---|---|---|---|
| Wild-type | Unmodified | none | 2.2 ± 0.51 |
| | H4K12/K16-acetylated | none | 0.35 ± 0.10 |
| | H4K12/K16-acetylated | 10 μM | 1.2 ± 0.20 |
| 4A | Unmodified | none | 7.3 ± 1.6 |
| | H4K12/K16-acetylated | none | 1.8 ± 0.22 |
| | H4K12/K16-acetylated | 10 μM | 3.5 ± 1.6 |

Means ± SE ($N$ = 3). $p300_{BRP}$, Wild-type of the BD-RING-PHD domain (residues 1048–1282) of human p300. 4A, the BD-RING-PHD domain (1048–1282) with mutations of R1133A, K1134A, R1137A, and K1140A.

**Supplementary Table 3 Dissociation constants between bromodomains (BD) and histone peptides measured by isothermal titration calorimetry**

| BD | Histone peptide (residues) | Modification | $K_D$ (μM) |
|---|---|---|---|
| $BRD4_{BD1}$ | H2B (1–27) | none | ND |
| | H2B (8–20) | K12ac/K15ac | 430 ± 5.0 |
| | H2B (16–27) | K20ac/K23ac | ND |
| | H4 (1–20) | none | ND |
| | H4 (1–20) | K5ac/K8ac | 49 ± 8.0 |
| | H4 (1–20) | K12ac/K16ac | 22 ± 7.0 |
| $p300_{BRP}$ | H2B (1–27) | none | ND |
| | H2B (8–20) | K12ac/K15ac | ND |
| | H2B (16–27) | K20ac/K23ac | 200 ± 30 |
| | H4 (1–20) | none | ND |
| | H4 (1–20) | K5ac/K8ac | 240 ± 8.5 |
| | H4 (1–20) | K12ac/K16ac | 15 ± 9.0 |

ND, not determined because $K_D$ is greater than 500 μM. Means ± SE. $BRD4_{BD1}$, the N-terminal BD (residues 44–168) of human BRD4. $p300_{BRP}$, the BD-RING-PHD domain (residues 1048–1282) of human p300.
